# Competing interactions give rise to two-state behavior and switch-like transitions in charge-rich intrinsically disordered proteins

**DOI:** 10.1101/2022.01.11.475920

**Authors:** Xiangze Zeng, Kiersten M. Ruff, Rohit V. Pappu

## Abstract

The most commonly occurring intrinsically disordered proteins (IDPs) are polyampholytes, which are defined by the duality of low net charge per residue and high fractions of charged residues. Recent experiments have uncovered nuances regarding sequence-ensemble relationships of model polyampholytic IDPs. These include differences in conformational preferences for sequences with lysine vs. arginine, and the suggestion that well-mixed sequences form a range of conformations, including globules, conformations with ensemble averages that are reminiscent of ideal chains, or self-avoiding walks. Here, we explain these observations by analyzing results from atomistic simulations. We find that polyampholytic IDPs generally sample two distinct stable states, namely globules and self-avoiding walks. Globules are favored by electrostatic attractions between oppositely charged residues, whereas self-avoiding walks are favored by favorable free energies of hydration of charged residues. We find sequence-specific temperatures of bistability at which globules and self-avoiding walks can coexist. At these temperatures, ensemble averages over coexisting states give rise to statistics that resemble ideal chains without there being an actual counterbalancing of intra-chain and chain-solvent interactions. At equivalent temperatures, arginine-rich sequences tilt the preference toward globular conformations whereas lysine-rich sequences tilt the preference toward self-avoiding walks. We also identify differences between aspartate and glutamate containing sequences, whereby the shorter aspartate sidechain engenders preferences for metastable, necklace-like conformations. Finally, although segregation of oppositely charged residues within the linear sequence maintains the overall two-state behavior, compact states are highly favored by such systems.

**Significance Statement:** Intrinsically disordered regions (IDRs) of proteins, when tethered to folded domains, function either as flexible tails or as linkers between domains. Most IDRs are polyampholytes that comprise a mixture of oppositely charged residues. Recent measurements of tethered polyampholytes showed that tendency of arginine- and lysine-rich sequences to behave very differently from one another. Using computer simulations, we show that these differences are determined by differences in free energies of hydration, steric volumes, and other considerations. Further, the interplay between electrostatic attractions and favorable free energies of hydration creates distinct stable states for polyampholytic IDRs. These findings have implications for switch-like transitions and the regulation of effective concentrations of interaction motifs by IDRs.

---

Significant fractions of eukaryotic proteomes are made up of intrinsically disordered regions (IDRs) (1). Conformational heterogeneity (2) is a defining hallmark of IDRs (3-5). Studies over the past decade have helped quantify relationships (6) that connect sequence-encoded information within IDRs to properties of conformational ensembles such as overall sizes and shapes, the amplitudes of spontaneous conformational fluctuations, and the dynamics of interconverting between distinct conformational states (7-19). These sequence-ensemble relationships have direct functional consequences that have been uncovered via studies based on biophysical, biochemical, and engineering approaches (5, 20-38). Our work, which is focused on physical principles underlying sequence-ensemble relationships of IDRs, is of direct relevance to understanding how IDRs function.

Charged residues are key determinants of sequence-ensemble relationships of IDRs (19, 39-41). They contribute through highly favorable free energies of hydration (42) and long-range electrostatic interactions. Net charge per residue (19, 39, 40) and the patterning of oppositely charged residues (43-45) are useful order parameters for describing sequence-ensemble relationships and interactions of charge-rich IDRs (46). Both features can be modulated through post-translational modifications (47-51), charge renormalization by solution ions (52), and charge regulation through context- and conformation-dependent uptake and release of protons (53).

Polyampholytes feature roughly equivalent numbers of oppositely charged residues, and they make up more than 70% of known IDRs (7, 17). For a given amino acid composition, which sets the fraction of charged residues and the net charge per residue, it has been shown that the linear mixing vs. segregation of oppositely charged residues can have a profound impact on sequence-ensemble relationships of polyampholytic IDRs (31, 32, 43). Specifically, for a given set of solution conditions, sequences featuring uniform linear distributions of oppositely charged residues are predicted to favor more expanded conformations compared to sequences with identical amino acid compositions where the oppositely charged residues are segregated into distinct blocks along the linear sequence. These predictions made using simulation and theory (43, 44, 54), have been confirmed using different experiments (31-33, 41, 55).

The ensemble-averaged radii of gyration (*R*_g_) of flexible polymers follow scaling relationships of the form *R*_g_ ∼ *N*^ν^. Here, *N* denotes the number of residues and the scaling exponent ν is a measure of the length-scale over which conformational fluctuations are correlated. For homopolymers or systems that are effective homopolymers, ν has four limiting values *viz*., 0.33, 0.5, 0.59, or 1, corresponding to globules, Flory random coils, self-avoiding walks, and rod-like conformations, respectively (56). Atomistic simulations performed at fixed temperatures suggest that ν ≈ 0.59 (43) for strong, well-mixed polyampholytes (17). The explanation for this behavior is as follows: Electrostatic attractions and repulsions are realized on similar length scales for well-mixed sequences. These interactions screen one another, and the highly favorable free energies of hydration become the main determinants of overall sizes and shapes of well-mixed strong polyampholytes (17). In contrast, compact conformations are formed by strong polyampholytes where oppositely charged residues are segregated into distinct blocks. Here, the electrostatic attractions between oppositely charged blocks can outcompete opposing effects of favorable solvation. These inferences were gleaned using sequences comprising 1:1 ratios of Lys and Glu (43). In the original simulations, the reference free energies of hydration of all charged residues were treated as being quantitatively equivalent and highly favorable. This leads to the hypothesis that Lys and Arg are interoperable with one another as determinants of sequence-ensemble relationships of IDRs (17). A similar inference emerges regarding the interoperability of Asp and Glu with respect to one another. The recent work of Sørensen and Kjaergaard has challenged these inferences (57). Using a system where model IDRs were deployed as flexible linkers between interaction domains, Sørensen and Kjaergaard used their measurements to estimate the relationships between amino acid sequence and the scaling exponent ν (57). Inferences from their experiments suggest that the ν ≈ 0.33 for (GRESRE)_*n*_ and ν ≈ 0.5 for (GKESKE)_*n*_ for the specific conditions they used in their measurements. Here, *n* is the number of repeats of the hexapeptides GRESRE or GKESKE. The results point to significant differences between Arg- and Lys-containing sequences. Further, while globularity of (GRESRE)_*n*_ has precedent in mean-field theories for polyampholytes, the mechanism by which Flory random coil-like behavior of (GKESKE)_*n*_ is achieved is unclear. Here, we develop a plausible physical explanation for the findings of Sørensen and Kjaergaard (57). Our work is based on atomistic simulations and the ABSINTH implicit solvation model and forcefield paradigm (58-61).

## Results

### Conformational ensembles of polyampholytic intrinsically disordered proteins (IDPs) show two-state behavior

Simulations were performed using an adaptation of recently recalibrated free energies of hydration (42, 62). These new values highlight the more hydrophobic nature of Arg when compared to Lys, and more favorable hydration of Asp / Glu when compared to Arg or Lys (see details in the *SI Appendix*). Using the recalibrated free energies of hydration (42, 62), we computed free energy profiles with *x* = (*R*_g_ / *N*^0.5^) as the reaction coordinate (see details in *SI Appendix*). Note that *x* ≡ *x*_FRC_ ≈ 2.5 Å per residue sets a useful reference length scale (63). Here, *x*_FRC_ is the value we obtain from our numerical approximation of the Flory random coil (FRC), which is an ideal chain model where all non-nearest neighbor interactions are ignored (2, 63). The free energy profile *W*(*x*) quantifies the free energy change associated with converting a Flory random coil to more expanded (*x* > *x*_FRC_) or compact (*x* < *x*_FRC_) conformations.

The free energy profiles, calculated at different simulation temperatures, are shown in Fig. 1A for (GKESKE)_7_. Seven repeats are short enough to be computationally tractable and long enough to observe the full spectrum of transitions without being confounded by finite size effects (43). The probability distribution function *P*(*x*), resolved along *x* as the reaction coordinate, is shown in Fig. 1B. Both *W*(*x*) and *P*(*x*) show that there are two stable states, one for *x* > *x*_FRC_ and another for *x* < *x*_FRC_ (*SI Appendix*, Fig. S3). In support of this two-state behavior, we note that the density of *P*(*x*) at *x* = *x*_FRC_ is essentially zero. The width of the well for *x* < *x*_FRC_ ranges from 1.4 to 2.2 Å per residue, and the minimum is located at *x* ≈ 1.6 Å per residue. Based on the scaling of internal distances, which shows plateauing behavior for low temperatures (*SI Appendix*, Fig. S2), we designate *x* ≈ 1.6 Å per residue, the location of one of the minima on the free energy profile, as *x*_globule_ (Fig. 1A). The width of the well for *x* > *x*_FRC_ ranges from 2.2 to 4.5 Å per residue. The minimum in *W*(*x*) and the corresponding peak in *P*(*x*) are located at *x ≈* 3.4 Å per residue. Scaling analysis shows that this free energy minimum is defined by a value of 0.6 for ν, implying that the minimum for *x* > *x*_FRC_ corresponds to a self-avoiding walk (*SI Appendix*, Fig. S2). Accordingly, we designate the minimum at *x* ≈ 3.4 Å per residue as *x*_SAW_ (Fig. 1A).

**Fig. 1.**
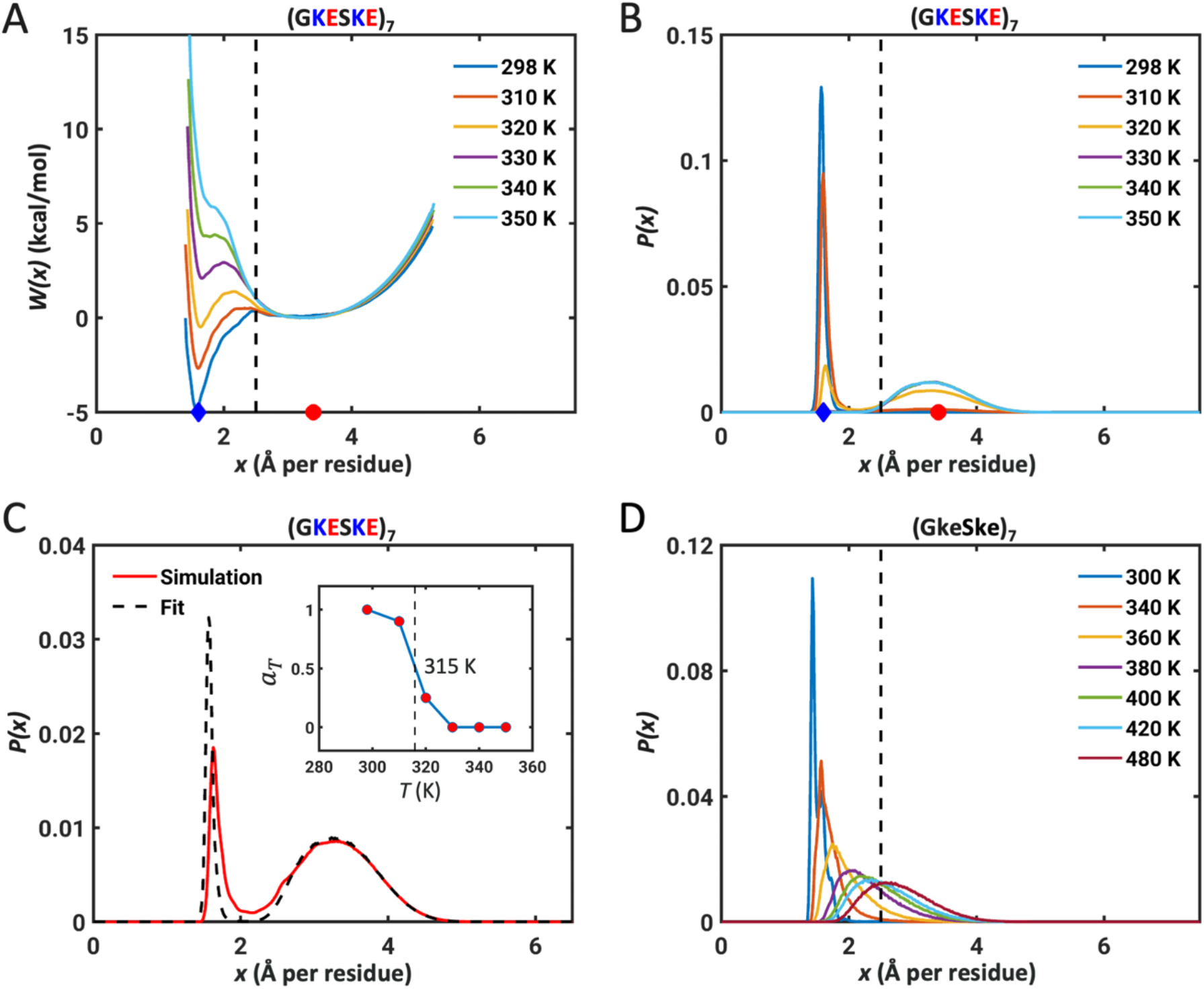
Neutral polyampholytic IDPs show two-state behavior. (A) The free energy profile *W*(*x*) for (GKESKE)_7_ calculated for different simulation temperatures. (B) Probability distribution functions *P*(*x*) obtained from the free energy profiles show the presence of two distinct peaks corresponding to the distinct minima in panel (A). The dashed line in (A) and (B) indicates the reference length scale of *x*_FRC_ = 2.5 Å per residue. And in each of the panels, the blue diamonds, and red circles mark positions of *x*_globule_ and *x*_SAW_, respectively. (C) The bimodal distribution *P*(*x*) can be fit to a two-state model (see main text). The fit is shown here for an intermediate simulation temperature of 320 K. The inset shows *a*_*T*_ as a function of temperature and the dashed vertical line corresponds to *T* = 315 K where *a*_*T*_ ≈ 0.5. (D) The distributions *P*(*x*) for the neutral polymer lacking charged residues change continuously as temperature increases. Note the existence of temperatures where distributions can be peaked around *x*_FRC_, which is not the case for the parent sequence (GKESKE)_7_ that has charged residues.

As temperature increases, the positions of the two minima in *W*(*x*) remain roughly fixed, while the relative depths, and the barrier separating the minima, change with temperature. At low temperatures, (GKESKE)_7_ favors globules. At high temperatures, the preferred state is the self-avoiding walk. At an intermediate temperature, globules and self-avoiding walks are of equivalent stability. The scaling exponent ν approaches 0.5 as the temperature of bistability is approached. To extract the temperature of bistability, we fit a two-state model to *P*(*x*) using a lever-rule: *P*(*x*;*T*) = *aT P*_globule_ (*x*)+ (1− *a*_*T*_) *P*_SAW_ (*x*). Here, 0 ≤ *a*_*T*_ ≤1; *P*_globule_(*x*) and *P*_SAW_(*x*) represent unimodal distributions peaked around *x*_globule_ and *x*_SAW_, respectively (Fig. 1C). These distributions were extracted from simulations at 298 K and 350 K, respectively. Using these reference distributions for each state, the estimated temperature of bistability, defined as the temperature where *a*_*T*_ = 0.5, is ≈ 315 K for (GKESKE)_7_ (inset in Fig. 1C).

We propose that the apparent FRC behavior reported by Sørensen and Kjaergaard (57) for the (GKESKE)_*n*_ system might derive from there being a bistability at the specific solution conditions that were investigated. Two-state behavior arises from competing interactions, namely, the highly favorable free energies of hydration of charged residues, which favor self-avoiding walks, and electrostatic attractions, which favor compaction. In contrast to charge-rich polyampholytic IDRs, the *P*(*x*) distributions for neutral polymers lacking charged groups should be continuous, tracking with the continuous change of the two-body interaction coefficient (6, 64). To demonstrate this, we computed temperature-dependent distributions for an artificial sequence designated as (GkeSke) _7_. Here, k and e are deprotonated and protonated versions of Lys and Glu, respectively. The free energies of hydration of deprotonated Lys and protonated Glu are ∼20 times lower in magnitude than the charged versions (*SI Appendix*, Table S2). Therefore, both competing interactions that are present in (GKESKE) _7_ are lost in (GkeSke)_7_, setting the two-body interactions due to short-range interactions as the main determinants of globule-to-coil transitions. As expected, the distributions for (GkeSke)_7_ show a continuous evolution from globules to self-avoiding walks, with there being at least one transition temperature where the distribution is peaked at *x*_FRC_ (Fig. 1D and *SI Appendix*, Fig. S4). Comparing the results in Fig. 1B to Fig. 1D suggests clear differences between the neutral, uncharged polymers vs. polyampholytes.

There is theoretical precedent for the observed two-state behavior of polyampholytes. For stiff homopolymers, the transition between globules and coils is sharp and the sharpness of this transition is governed by the interplay between the two- and three-body interactions (65-67). Kundagrami and Muthukumar (68) introduced a three-body interaction term to generalize an earlier theory put forth by Muthukumar (69) to demonstrate that a polyelectrolyte can undergo a first-order coil-to-globule transition. The existent of a bistability is tied to a discontinuous change in the effective charge that is due a charge regularization enabled by adsorbed counterions (68). Interestingly, the discontinuity of effective charge was associated with dielectric inhomogeneities around the polymer backbone. This is a defining feature of the ABSINTH model that engenders competing interactions.

Recently, Ghosh and coworkers have extended the approach of Muthukumar to describe the collapse transitions of finite-sized heteropolymers. They report the existence of an effective temperature of bistability that results from the choice they make for the free energy functional for polyampholytic systems (54, 70). Using a mean-field description for random polyampholytes, an approach that ignores chain connectivity and sequence details, Higgs and Joanny showed that in the long-chain limit of *N* → ∞, individual random polyampholytes collapse to form globules irrespective of the sign or magnitude of the two-body interaction coefficient (71). The globular state is favored by electrostatic attractions. In contrast to the long-chain limit, finite-sized chains in the theory of Higgs and Joanny are predicted to form expanded conformations that are akin to self-avoiding walks (71). The preference for collapsed states (72, 73), whereby electrostatic attractions drive chain collapse, was also predicted by the Flory theory for individual chains of random polyampholytes that was developed by Dobrynin and Rubinstein (74). Our observations of two-state behavior mimic those of Moldakarimov et al., who noted that neutral polyampholytes undergo jump-like coil-to-globule transitions due to the formation of intra-chain ion pairs (75). The two-state behavior and the nature of the collapse transition can be modulated by an excess of charge of one type (76-78).

The main finding is that a scaling exponent of 0.5 can arise because the conformations are mixtures of globules, defined by a negative excluded volume, and SAWs, defined by a positive excluded volume. In contrast, observations of scaling exponents of 0.5 are typically taken to mean that the polymer in question is at its theta temperature with an excluded volume of zero. The challenge is to discern between these two scenarios, both of which yield an apparent scaling exponent of 0.5. This requires the deployment of measurements that directly access or allow one to infer the distributions of polymer sizes. This seems feasible using single molecule measurements that access distributions of radii of gyration or internal distances (39, 79, 80). Advances in single molecule detection might make such experiments feasible (81). A single molecule variant of the measurements performed by Sørensen and Kjaergaard could be deployed to uncover the distributions of effective concentrations. The measurements will have to be of coil-to-globule transitions as a function of a suitable perturbant. Unlike the continuous transitions that have been reported for most systems, the expectation is of fixed two-state behavior (if temperature is the perturbant) or variable two-state behavior (if denaturant is the perturbant) (82). The signatures of bistable vs. continuous transitions can be gleaned by quantifying the distributions of *R*_g_ and effective concentration values at the system-specific apparent theta temperatures. This is shown in **Fig. 2A** and **Fig. 2B**. Whether or not the presence of a bistability can be detected using single molecule measurements will likely depend on the timescales for the transitions between globule and SAW states.

**Fig. 2:**
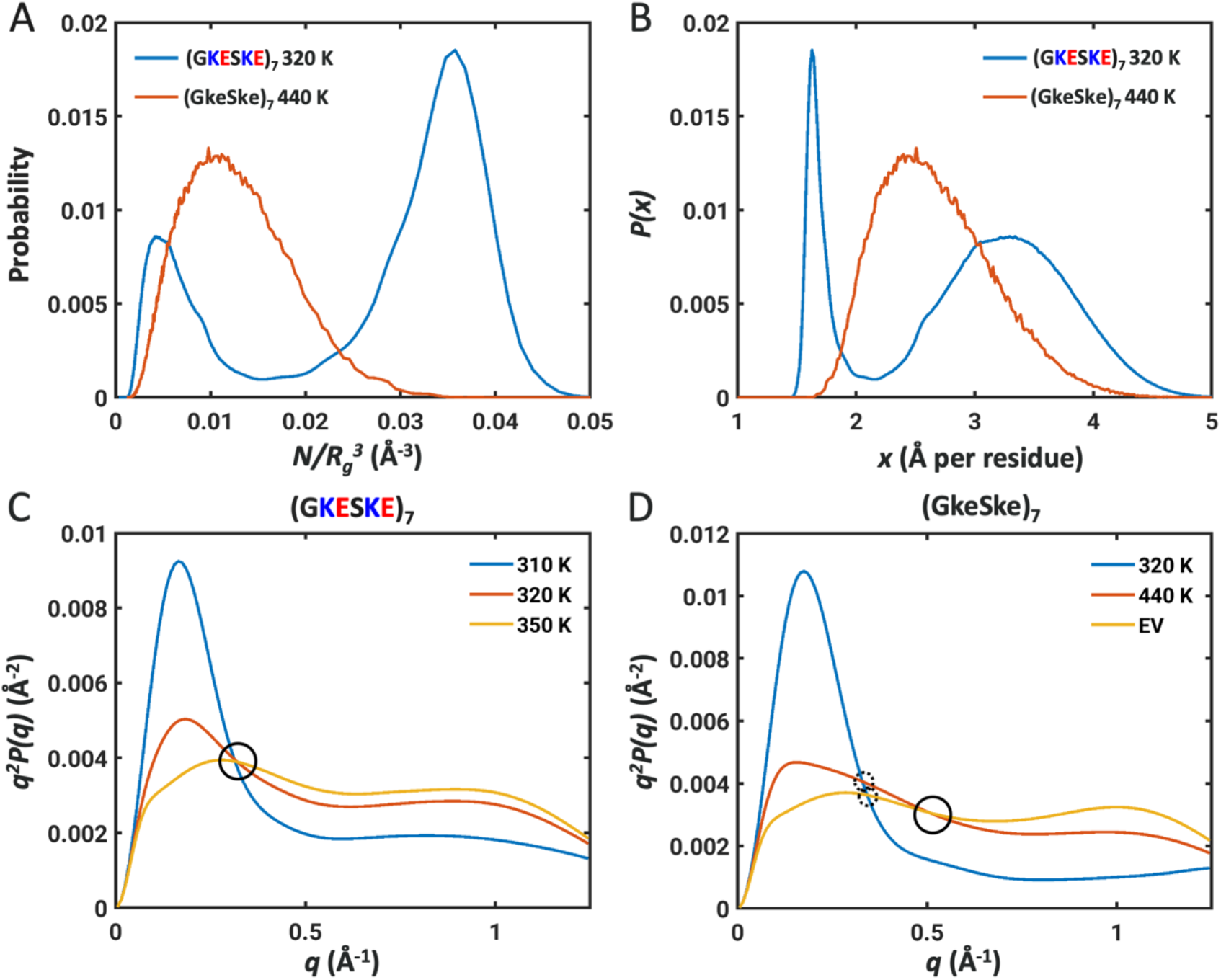
*R*_*g*_distributions and Kratky profiles at the apparent theta temperature for two types of systems where the scaling exponents would be 0.5. (A) Differences in *R*_*g*_ distributions, plotted as distributions of effective concentrations at the apparent theta temperatures for (GKESKE)_7_ (blue) vs. (GkeSke)_7_ (red). (B) Differences in distributions of *x*, the normalized *R*_*g*_ distributions between the two systems at their respective apparent theta temperatures. The bimodality of a bistable system is replaced by a Gaussian behavior for system that lacks charged residues. (C) Kratky profiles for globules (blue), coils (red), and yellow (SAWs) for (GKESKE)_7_ computed at the relevant temperatures. The single crossover length scale is circled for ease of identification. (D) Kratky profiles for globules (blue), coils (red), and yellow (SAWs) for (GKESKE)_7_ computed at the relevant temperatures. Non-coincidence of crossover length scales is shown as dashed circles, and the presence of an additional crossover length scale is identified using a solid circle.

Can differences between the two types of behaviors be detected using an ensemble measurement such as small angle x-ray scattering (SAXS)? To answer this question, we computed Kratky profiles (83) for (GKESKE)_7_ vs. (GkeSke)_7_. For each system, three specific temperatures were chosen, and these correspond to temperatures where the systems of interest are characterized by apparent scaling exponents (ν_app_) of ≈ 0.33, ≈ 0.5, and ≈ 0.59. For the (GKESKE)_7_ system we observe Kratky profiles with two distinct peaks located at similar positions, and the three Kratky profiles crossover at a common wavenumber. The presence of a single crossover length scale, circled in **Fig. 2C**, as a function of temperature or other perturbants is to be contrasted with the Kratky profiles for the (GkeSke)_7_ system (**Fig. 2D**). For the system without charged residues, the continuous nature of the globule-to-SAW transitions give rise to non-coincident cross-over length scales for the globule and coil vs. globule and SAW, plus an additional crossover between the profiles for coils and SAWs.

### Temperature of bistability and the nature of the collapse transition are affected by sidechain chemistries

The two-state behavior and the presence of a temperature of bistability arises from the competing effects of electrostatic attractions and favorable free energies of hydration. Therefore, we propose that the sidechain specific reference free energies of hydration, which arise from differences in chemical structures of sidechains, combined with chemistry-specific interactions of sidechains should affect the temperature of bistability and the nature of the collapse transition. We tested this hypothesis using ABSINTH-based simulations of (GRESRE)_7_, (GKDSKD)_7_, and (GRDSRD)_7_. Being able to capture distinct preferences of Arg-vs. Lys-containing sequences requires a formal accounting of the differences in free energies of hydration that were derived recently using a combination of experimentally derived quantities and free energy calculations (42, 62). These results indicate that despite being a strong base, arginine is more hydrophobic than lysine, and the physical basis for this was discussed recently by Fossat et al (42).

We obtained free energy profiles and probability density functions, *W*(*x*) and *P*(*x*), for (GRESRE)_7_, (GKDSKD)_7_, and (GRDSRD)_7_ as shown in *SI Appendix*, Fig. S6. For these simulations, we used recalibrated free energies of hydration that are summarized in *SI Appendix*, Table S2. The qualitative behaviors of all three systems resemble those of (GKESKE)_7_. These results suggest that the two-state behavior is a generic attribute of well-mixed polyampholytes. However, there are quantitative differences among the systems. These are summarized in **Fig. 3**, where we plot the ensemble average of *x* against reduced simulation temperatures for (GKESKE)_7_, (GRESRE)_7_, (GKDSKD)_7_, and (GRDSRD)_7_.

**Fig. 3.**
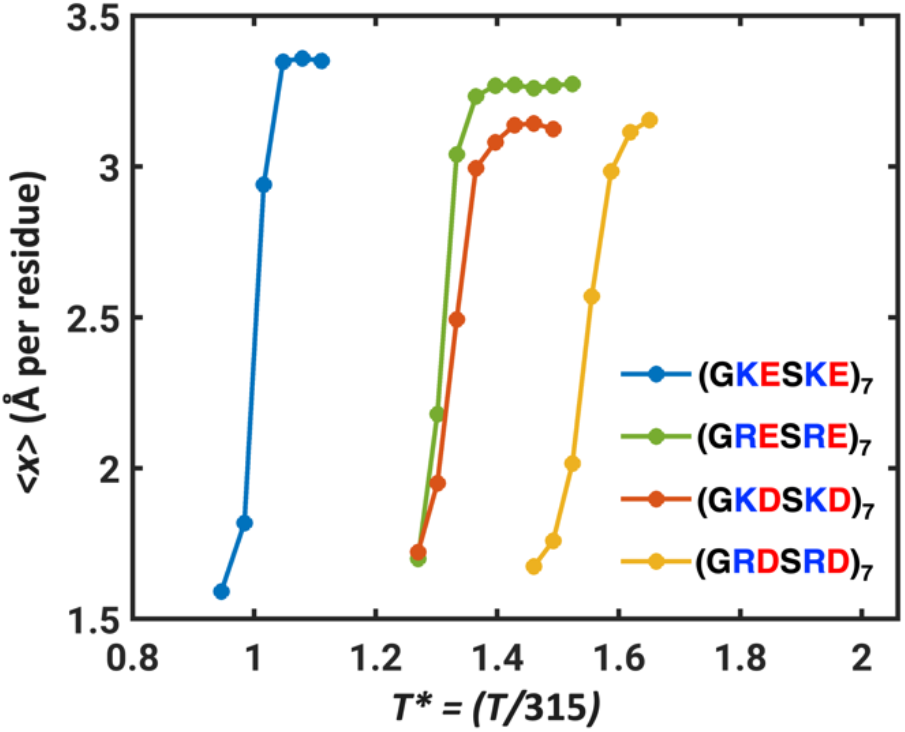
Ensemble-averaged values of *x*, the normalized *R*_g_, plotted against simulation temperature for (GKESKE)_7_, (GRESRE)_7_, (GKDSKD)_7_, and (GRDSRD)_7_.

The reduced temperature is *T*\* = (*T* / 315 K) where *T* is the simulation temperature, and 315 K corresponds to the inferred simulation temperature of bistability for (GKESKE)_7_. The temperatures in our simulations are not to be taken literally given the mean-field nature of the solvation model. Instead, interpretation of simulation temperatures in terms of realistic solution conditions will require some prior calibration against experimental data for the system of interest – as has been shown in recent work, albeit in different contexts (46, 84). Comparisons in terms of the reduced temperature *T\** indicate that replacing Lys with Arg shifts the temperature of bistability upward by a factor of ∼1.3 for (GRESRE)_7_. Likewise, the temperature of bistability is higher by a factor of ∼1.2 for (GRDSRD)_7_ compared to (GKDSKD)_7_. It follows that Arg as the basic residue tilts the balance toward globules when compared to Lys, which tilts the balance toward self-avoiding walks (*SI Appendix*, Fig. S8).

In addition to differences between Lys and Arg, we find that replacing Glu with Asp also has an impact on the transition between globules and self-avoiding walks. Specifically, the apparent temperature of bistability is higher by a factor of ∼1.3 for (GKDSKD)_7_ when compared to (GKESKE)_7_. Likewise, the apparent temperature of bistability is higher by a factor of ∼ 1.25 for (GRDSRD)_7_ compared to (GRESRE)_7_. These differences cannot be explained in terms of differences in free energies of hydration (see *SI Appendix*, Table S2). Instead, we considered two plausible explanations for the differences we observe: First, the longer sidechain of Glu and the higher sidechain entropy could lower the temperature of bistability. Second, the longer sidechain of Glu and the higher steric volume will likely weaken electrostatic attractions within the interior of globules.

To test the two hypotheses, we compared profiles for temperature-dependent collapse transitions of two artificial sequences (GkeSke)_7_ and (GkdSkd)_7_. Here, k is the deprotonated Lys, whereas e and d are protonated versions of Glu and Asp, respectively. If differences in sidechain entropy are the main contributors, then it should follow that (GkdSkd)_7_ will have a higher transition temperature than (GkeSke)_7_. However, as shown in *SI Appendix* Fig. S9, the transition temperature for (GkdSkd) _7_ is lower than that of (GkeSke)_7_, thereby ruling out the first hypothesis. We tested the second hypothesis by comparing the transition temperature of (GKESKE)_7_ to that of (GZESZE)_7_ where Z is 2,4-diamino butyric acid or Dab. This maintains the charge and chemical structure of the amine while replacing the longer sidechain in Lys with the shorter sidechain of Dab. As shown in *SI Appendix*, Fig. S10, the transition temperature for (GZESZE)_7_ is ∼1.3 times higher than (GKESKE)_7._ This result implies that charged residues with equivalent functional groups and shorter sidechains stabilize compact conformations when compared to sequences with charged residues that have longer sidechains. Accordingly, we ascribe the differences between (GRDSRD)_7_ and (GRESRE)_7_ to the fact that the longer sidechain of Glu and its higher steric volume weaken electrostatic attractions within the interior of globules.

To further interrogate the differences between different well-mixed polyampholytes, we computed free energy profiles and distribution functions for sequences that are essentially polyzwitterions (86). These include (RE)_25,_ (KE)_25_, (RD)_25_, and (KD)_25_ (Figs. 4A-B and *SI Appendix*, Figs. S11, S12). Our investigations of these systems were motivated by recent work that show the relevance of RD and RE repeats for nuclear speckle assembly (87). This work also highlighted profound differences in the phase behavior of Arg vs. Lys containing polyzwitterions. The Glu-containing polyzwitterions show two-state behavior, whereas the Asp-containing sequences show the presence of an intermediate state that is favorably populated at the transition temperature. We examined two-dimensional histograms *p*(*x, u*_ABSINTH_) computed in terms of *x* and the conformation-specific values of the ABSINTH potential energies. These histograms confirm the two-state behavior of (RE)_25_ (Fig. 4C). However, for (RD)_25_, we observe an intermediate state between globules and self-avoiding walks (Fig. 4D). These intermediate states correspond to “necklace-like” conformations (Fig. 4G) that have been predicted to be either stable or metastable states by different theories for polyampholytes (76). We compute differences in the numbers of hydrogen bonds formed by Asp containing polyzwitterions. These differences are most pronounced for conformations with values of *R*_g_ that are on a par with or larger than Flory random coils (*SI Appendix*, Fig. S13). The Asp containing sequences feature more hydrogen bonds, in expanded conformations, when compared to Glu containing sequences. Accordingly, it appears that necklace-like conformations formed (RD) and even (KD) repeats are stabilized by hydrogen bonds enabled by the shorter Asp sidechain.

**Fig. 4.**
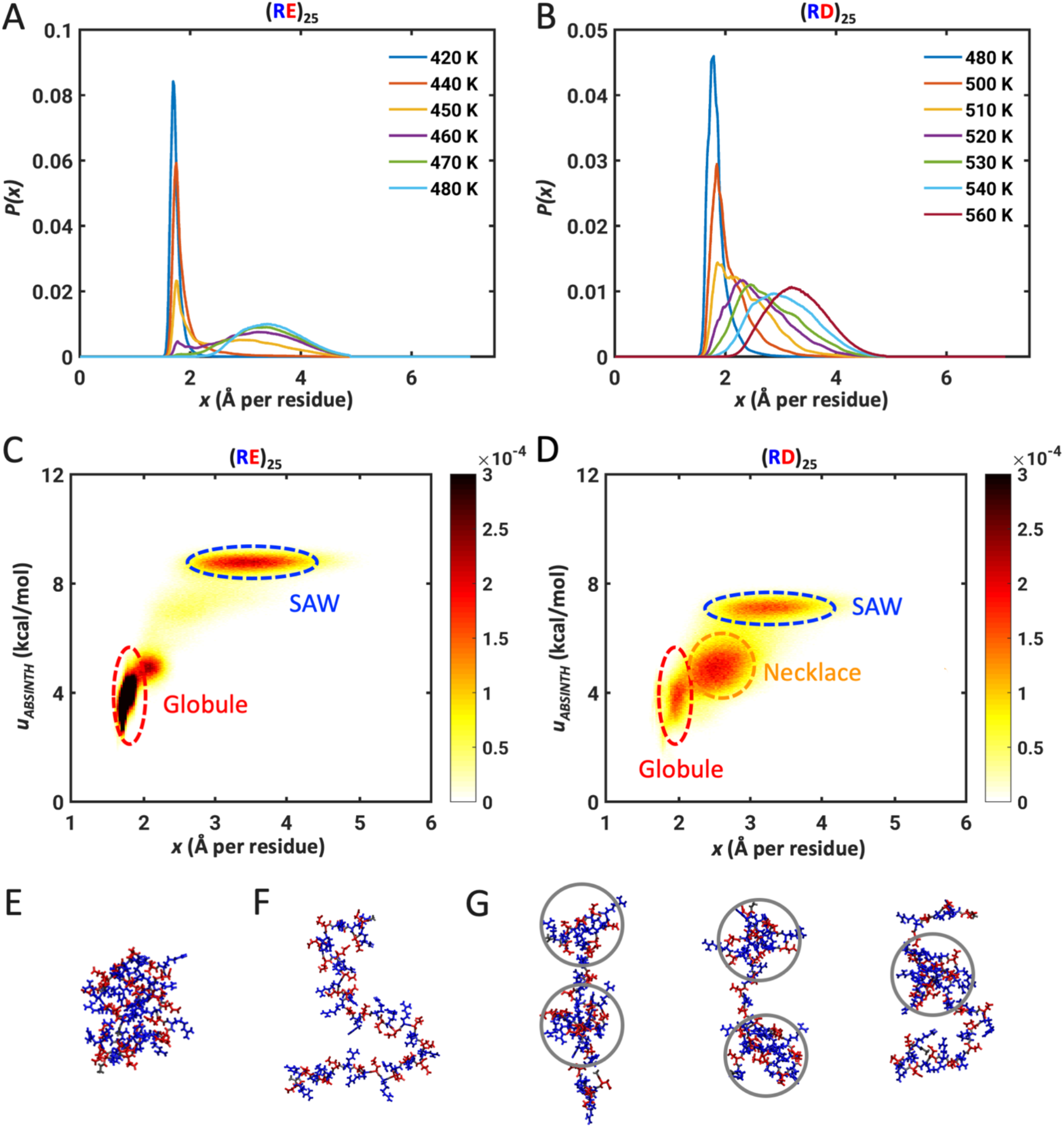
Asp residues modulate the two-state behavior of polyampholytic IDPs. Calculated distributions *P*(*x*) for (A) (RE)_25_ and (B) (RD)_25_. Two-dimensional histograms *p*(*x, u*_ABSINTH_) for (C) (RE)_25_ and (D) (RD)_25_. Here, *u*_ABSINTH_ is relative potential energy per residue, referenced to the system-specific lowest energy conformation. The histograms were computed as an average over unbiased replica exchange simulations at three temperatures near the transition temperature, namely 450 K, 460 K and 470 K for (RE)_25_, 520 K, 530 K and 540 K for (RD)_25._ The bin widths for *x* and *u*_ABSINTH_ are 0.004 Å and 0.02 kcal/mol per residue, respectively. (E) Snapshot of a representative globular conformation for (RE)_25_. (F) Snapshot of a representative self-avoiding-walk conformation for (RE)_25_. (G) Three snapshots, showing representative necklace-like conformations, with gray circles delineating the “pearls” formed by (RD)_25_. Structures were drawn using VMD (85). Lys is shown in blue and Glu / Asp residues are in red.

### Influence of sequence patterning on the two-state behavior of polyampholytic IDRs

Next, we investigated the impact of the linear segregation of oppositely charged residues on the two-state behavior of polyampholytes. For this, we performed simulations for two Lys and Glu containing sequences designated as sv5 and sv10 (**Fig. 5A**) (43). The results in **Figs. 5 B-C** show that these two sequences also exhibit two-state behavior. However, the peak in the distribution *P*(*x*) at high temperatures is much smaller (∼2.7 Å per residue) for sv10 when compared to that of sv5 (3.6 Å). For self-avoiding walk-like conformations, the value of *x* should have a peak around 3.4 Å per residue. This is not the case for sv10. Instead, the distribution *P*(*x*) at high temperature for sv10 has a wide right shoulder that can be fit to a mixture of two Gaussian distributions, one peaked at 2.7 Å per residue, and the other peaked at 3.2 Å per residue. This implies that the blockier sequence populates a metastable state, which for sv10 corresponds to Ω-loop-like structures (88) that are shown in **Fig. 5D**. These metastable structures are stabilized by electrostatic attractions between the N-terminal Lys patch and C-terminal Glu patch. The close contacts between N-terminal and C-terminal are clearly shown in the distance and scaling maps (*SI Appendix*, Fig. S16). Overall, increased charge segregation increases the stability of different compact structures that are globule-, loop-, and hairpin-like (*SI Appendix*, Fig. S17).

**Fig. 5.**
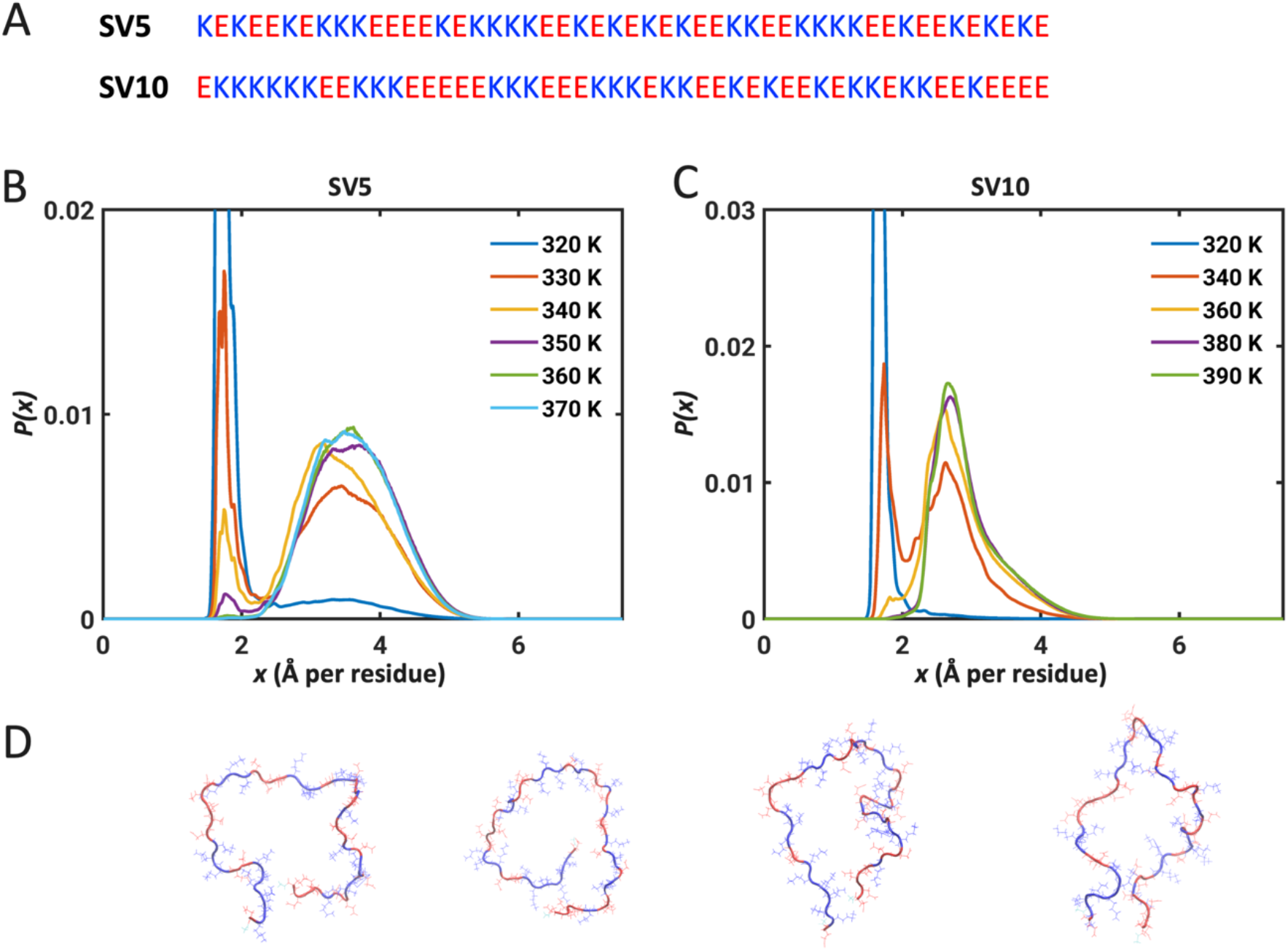
Two-state behavior is preserved for sequences with blocks of oppositely charged residues. (A) Amino acid sequences of sv5 and sv10 – nomenclature as in Das and Pappu (43). Panels (B) and (C) show the distributions *P*(*x*) computed for two sequences designated as sv5 and sv10. The corresponding free energy profiles of (B) and (C) are shown in *SI Appendix* Fig. S14 (B) *P*(*x*) for sv5 shows that the two-state behavior is preserved for sequences with mild degrees of segregation of oppositely charged residues. (C) *P*(*x*) for sv10 shows that there are temperatures where intermediate states, corresponding to well-defined metastable structures, become prominent as the linear segregation of oppositely charged residues increases. (D) Four Ω-loop-like structures (88) observed as stable intermediates for sv10. The structures were extracted from simulations performed at 340 K and were drawn using VMD (85). Lys is in blue and Glu residues are in red.

### Dynamics of interconversion between globules and self-avoiding walks

To investigate the dynamics of interconversion between globules and self-avoiding walks, we performed Langevin dynamics simulations using the free energy profiles extracted for (GKESKE)_7_ (see details in *SI Appendix*). These simulations were performed using three different profiles (Fig. 6A), one corresponding to the case where the globule is more stable than the self-avoiding walk (*K*=8.7), the second corresponding to the case of bistability (*K*=1.0), and the third corresponding to the case where the self-avoiding walk is more stable than the globule (*K*=0.3). From the simulation trajectories, we calculated the mean residence times to the left and right of the dividing line, depicted as a dotted line in Fig. 6A. The mean residence times track with the equilibrium constants, being longer in the globule region (left of the barrier) for *K =* 8.7 and longer in the coil region (right of the barrier) for *K* = 0.3 (Fig. 6B). In addition to the mean residence times (*t*_res_), we computed the mean time for transitioning from the globule-to-coil basins and vice versa. To account for the asymmetries of the globule and coil basins, we computed the basin-to-basin transit times (τ_basin_) as the time it takes for transitioning from the window striped in red to that in blue and vice versa. Given that a barrier must be negotiated for these transitions to occur, we find that τ_basin_ is greater than *t*res by an order of magnitude (Fig. 6C). For our analysis of τbasin and *t*res we used *Rg* as the reaction coordinate. This is relevant because the transitions between globules and self-avoiding walks are associated with the change in internal density and large fluctuations in density around the barrier (delineated in Fig. 6A) as noted by Kuznetsov and Grosberg (89). It is worth emphasizing that estimation of τbasin rests on the choice of two regions of equal width within the globular and coil regions. However, the potential of mean force is an asymmetrical double well, with the coil basin being broad and shallow when compared to the globule basin – even at the temperature of bistability. In such systems, the mean first passage time for crossing the barrier will be influenced by the diffusive search within the broad and shallow basin, thereby yielding an effective kinetic readout that might confound simple interpretations of two-state behavior (90).

**Fig. 6.**
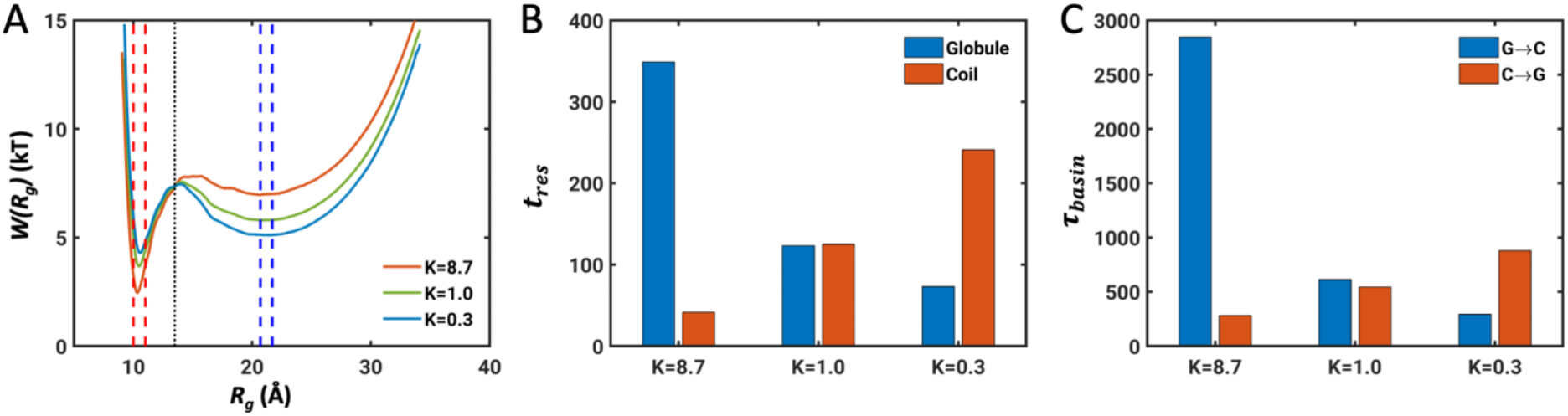
Simulated dynamics for transitions between globules and self-avoiding walks. (A) Free energy profiles *W*(*R*_*g*_) used for Langevin dynamics simulations. The dashed line indicates the boundary between the globule and coil state. Regions of equal width within the globule and SAW basins are indicated by the red and blue dashed windows, respectively. (B) Mean residence times (*t*_res_) within the globule and coil (self-avoiding walk) states. (C) Mean basin-to-basin transit times (τ_basin_) for forward and reverse transitions between globule (red region in panel A) to the coil (SAW) basins. Details of how we calculated τ_basin_ are described in the *SI Appendix*.

## Discussion

In this work, we have uncovered the two-state behavior of strong polyampholytic IDPs. At a given temperature, the relative stabilities of the two states are modulated by the identity of the basic residue, Lys vs. Arg (Fig. 2). Our findings provide a plausible explanation for the observations of Sørensen and Kjaergaard (57). We show that the apparent Flory random coil-like scaling behavior reported for the (GKESKE)n system arises from an averaging over the contributions of coexisting globules and self-avoiding walks. Further, we find that the two-state behavior is altered to accommodate metastable necklace-like, conformations when the acidic residue is Asp as opposed to Glu. Although the two-state behavior is preserved for polyampholytes with blocky architectures where oppositely charged residues are segregated from one another, compact, globule-, loop-, and hairpin-like conformations are highly stable due to intra-chain electrostatic attractions (*SI Appendix*, Fig. S17). Detection of the proposed bistable behavior will require experiments that use temperature, salt, denaturant, or pH as perturbants. Ideally, the experiments will be able to uncover population distributions, not just ensemble averages, although in systems with high fractions of charged residues even ensemble averaged measurements might be valuable (Fig. 2).

Our findings establish that competing effects of favorable hydration of charged groups, differences in steric volumes (*SI Appendix*, Fig. S8), and intra-chain electrostatic interactions among oppositely charged residues give rise to two distinct states namely, self-avoiding walks and globules. The relative stabilities are influenced by: (i) temperature, which is a proxy for solvent quality; (ii) the identity of the basic residues, with Arg favoring globules more so than Lys; (iii) the linear segregation vs. mixing of oppositely charged residues, where increased segregation gives rise to stronger preferences for compact states; (iv) the fraction of charged residues and the net charge per residue, with lower fractions of charged residues enabling conventional, continuous coil-to-globule transitions, and increased charge of one kind driving the preference for self-avoiding walk-like conformations (19, 39, 40). The differences between Arg and Lys as well as the differences between Asp and Glu (albeit to a lesser extent) have received recent attention in the context of work on the molecular grammar underlying the driving forces for phase separation of IDPs (46, 87, 91-93). As noted by Tesei et al., (94) different hydrophobicity scales show a lack of consensus regarding the apparent hydrophobicity of Arg vs. Lys. Arriving at a consensus will require the assessment of differences in intrinsic differences in free energies of hydration and in the context dependent effects of Arg vs. Lys and Glu vs. Asp (42, 62). Additionally, the sensitivity of a sequence to the effects of charge regulation will also be highly dependent on the identities of ionizable residues as well as sequence contexts (53, 95).

Metastable, necklace-like conformations are accessible to sequences featuring differences in the lengths of sidechains of acidic vs. basic residues. Examples of such systems include RD repeats that are found in IDRs within nuclear speckle proteins (87). RD repeats were shown to form spherulitic fibrils through intermolecular associations, and the formation of fibrillar solids is governed by the number of RD repeats and overall charge neutrality (87). We propose that necklace-like conformations are likely to be the drivers of intermolecular associations that give rise to fibrillar structures. This hypothesis is based on the observation that the necklace conformation might be an excited state corresponding to the so-called N* state proposed by Thirumalai and coworkers (96, 97) as being essential for driving fibril formation. Greig et al., (87) also showed that Asp and Glu play different roles in the nuclear speckle condensation. Whereas RD repeats form fibrils, the RE repeats form non-fibrillar condensates. Further, the driving forces for assembly are stronger for the RD repeats when compared to RE repeats. Our studies establish clear differences in the free energy landscapes of these systems. It remains to be ascertained if these differences help explain the finding of Greig et al. (87).

For infinitely long homopolymers or effective homopolymers the scaling exponent ν has four limiting values *viz*., 0.33, 0.5, 0.59, or 1. These values correspond to globules, Flory random coils, self-avoiding walks, and rod-like conformations, respectively (56). Deviations from these exponents are expected for finite sized homopolymers, and the magnitude of finite size corrections that are required can be formally estimated (98, 99). Unlike homopolymers, unfolded proteins and IDPs are complex, finite sized, heteropolymers. Such systems feature a spectrum of intra-chain and chain-solvent interactions. Accordingly, a value for ν_app_ that is not 0.33 or 1 could arise from a mixture of stable states – a point that is made in this work for strong polyampholytes. Our findings highlight the importance of going beyond estimates for ν_app_ as a device for comparative assessments of sequence-ensemble relationships of unfolded proteins and IDPs. What we need to measure are distributions for the order parameter, and this seems feasible using modern single molecular spectroscopies.

The two-state behavior of polyampholytic IDPs allows for the formal possibility of switch-like transitions between globules and self-avoiding walks. These transitions can be spontaneous if the solution conditions are such that the system of interest is in the vicinity of the temperature of bistability. Away from this temperature, switch-like transitions can be driven by energy inputs or post-translational modifications. Overall, the dynamics of interconversion between stable states will be influenced by sequence-intrinsic properties of the free energy landscapes such as the barrier for interconversion and the widths of conformational basins around the stable states (Fig. 6).

Scanning for solution conditions (79, 100, 101) that place polyampholytic IDPs in the vicinity of the temperature of bistability could be a way to uncover the prospect of switch-like interconversions between dramatically different conformational states of equivalent stability. Precedent for switch-like transitions and dynamics that span a spectrum of timescales has been reported for different IDRs (10, 13). Such transitions, especially in the context of tethered systems, are likely to make important contributions to the spatial and temporal control over biochemical reactions where IDRs modulate the effective concentrations of ligands around binding sites or of substrates around active sites of enzymes (31, 32, 34, 57, 102-106). While our results are presented for IDRs studied as autonomous units, we expect that our findings will be transferable to tethered IDRs, albeit with context-dependent modifications (104, 107).

The simulations reported here were performed in the absence of excess salt or solution ions (if the systems were electroneutral, see *SI Appendix*). To probe the effects of excess salt, we investigated the effects of adding 50 mM NaCl for the (GRESRE)_7_ system. The results are summarized in *SI Appendix* Fig. S7. The temperature of bistability shifts down in the presence of salt, and this behavior, which is consistent with the observations of Kundagrami and Muthukumar (68) for polyelectrolutes, can be rationalized as being mainly due to the screening effects of monovalent salts. Solutions ions are likely to play an important role in affecting conformational transitions, conformational equilibria, binding equilibria, and phase equilibria of polyampholytic and polyelectrolytic IDRs (24, 39, 52, 79, 91, 108, 109). Besides the screening effect, predicted by Higgs and Joanny (71) and illustrated in *SI Appendix* Fig. S7, there is the expectation that ions that preferentially accumulate around blocks of the same type of charge are released when the ion-chain interactions are replaced by intra-chain interactions (68, 69, 109). The effects of solution ions are also likely to be directly relevant to the phase behaviors of polyampholytic IDRs (110, 111).

Finally, we asked if sequences that are likely to show bistability are present in the disordered proteome. For this, we calculated the distribution of fraction of charged residues (FCR) for IDRs in DisProt 2022-03 database (*SI Appendix*, Fig. S18). We also calculated FCR values of strong polyampholytes (17), and the fractions of specific charged residues. This analysis uncovered 202 out of 3236 IDRs with lengths ≥ 30 and FCR values equal to or larger than 0.5. Further, 26 out of 299 strong polyampholytes with lengths ≥ 30 have FCR values equal or larger than 0.5. Asp disfavors the bi-stable distribution, but it has a lower frequency of being observed when compared to Glu. Higher fractions of Lys when compared to Arg also indicate that strong polyampholytes are likely to show a preference for bistability since Arg rich polyampholytes tend to prefer globules. *SI Appendix* Table S6 lists sequences for the top 15 strong polyampholytes with length ≥ 30 ranked by FCR in the DisProt 2022-03 database. Annotations of functions suggest that some of the IDRs are thought to function as flexible linkers that are regulated by multisite phosphorylation.

## Materials and Methods

### Monte Carlo simulations

All simulations were performed using version 2.0 of the CAMPARI molecular modeling software (http://campari.sourceforge.net/). The forcefield parameters were derived based on OPLS-AA/L forcefield as implemented in abs3.2_opls.prm. Free energies of hydration for the charged residues were based on recently recalibrated values. Na^+^ or Cl^-^ ions (112) were added to the simulation droplet to neutralize the system if the net charge of the peptide is not zero. All the details are described in *SI Appendix*.

### Free energy profiles

Umbrella sampling was used to obtain free energy profiles using *x* as the reaction coordinate. Thermal replica exchange Monte Carlo simulations (113) were used to enhance the sampling of globular structures in the windows with low *R*_g_. Standard Metropolis Monte Carlo simulations were performed at the rest of windows. The weighted histogram analysis method (114) was used to derive the free energy profile. The details of the setup for each system, the umbrella sampling, replica exchange, and reweighting are described in *SI Appendix*.

## Acknowledgments

This work was supported by grants from the US National Institutes of Health (R01NS121114 and 5R01NS056114) and the US Air Force Office of Scientific Research (FA9550-20-1-0241). We thank Samuel Cohen, Martin Fossat, Alex Holehouse, Greg Jedd, Ammon Posey, Min Kyung Shinn, and Andrea Soranno for useful discussions regarding charge-rich IDRs. XZ would like to thank Louis Smith for helpful discussions and Jared Lalmansingh, Stephen Tahan and Mark Bober for support in the using McKelvey Engineering Compute Cluster and RIS cluster at Washington University in St. Louis.

