## Supporting Information Appendix for "Competing interactions give rise to two-state behavior and switch-like transitions in charge-rich intrinsically disordered proteins"

for

Xiangze Zeng, Kiersten M. Ruff and Rohit V. Pappu[1]
Department of Biomedical Engineering and Center for Science & Engineering of Living Systems, Washington University in St. Louis, St. Louis, MO 63130

### 1. Details of Metropolis Monte Carlo (MC) simulations

Metropolis Monte Carlo simulations for the were performed using version 2.0 of the CAMPARI molecular modeling software (http://campari.sourceforge.net/). The simulations used the ABSINTH implicit solvent model and underlying forcefield paradigm (1). The forcefield parameters were derived based on OPLS-AA/L forcefield (2) and implemented in the abs3.2_opls.prm parameter set. Parameters for proline residues were based on the work of Radhakrishnan et al (3). The values of reference free energies of hydration for the side chains of the charged amino acids, $Arg^{+}$, $Lys^{+}$, $Asp^{-}$ and $Glu^{-}$ were calculated using the TCPD method of Fossat et al (4) with -252.7 kcal/mol as the reference free energy of hydration of the proton. The input parameters for the TCPD method are listed in Table S1. The values of the free energy of hydration from the TCPD method were further calibrated and shifted by -11.5 kcal/mol so that the ensemble averaged $R_g$ $(GKESKE)_{10}$ at 298 K was close to the reference length scale $x_{FRC} = 2.5$ Å of the Flory random coil and that the scaling exponent, estimated from the scaling of internal distances, was near 0.5 as reported by Sørensen and Kjaergaard (5). Temperature-independent reference free energies of hydration were used in this study. Table S2 lists the values used for the sidechains of charged residues. The temperature-dependent macroscopic dielectric constant of water was set based on previous work (6). The cutoff for the short-range Lennard-Jones potential was set to be 10 Å. The cutoff for electrostatic interactions between different sites on neutral groups was 14 Å. There were no cutoffs for electrostatic interactions involving mobile ions and charged side chains of $Arg^{+}$, $Lys^{+}$, $Asp^{-}$ and $Glu^{-}$.

The polyampholytic IDPs were simulated in a spherical droplet with the radius of 100 Å. Randomly generated self-avoiding conformations were used as the initial conformations for the simulations. Move sets for Metropolis Monte Carlo (MC) simulations included side-chain torsions, concerted rotations, pivot moves, translations of ions, and translations and rotations of individual molecules. For thermal replica exchange Monte Carlo (REMC) simulations, swaps between neighboring replicas were proposed once every $5 \times 10^4$ MC simulation steps.

For unbiased simulations of $(GkeSke)_7$ and $(GkdSkd)_7$, two independent REMC simulations were performed for each system. Each independent simulation consisted of $10^8$ MC steps at one temperature, and the temperature was incremented from 260 K to 480 K in steps of 10 K. For the unbiased REMC simulations of $(RE)_{25}$ and $(RD)_{25}$, one long simulation consisting of $4 \times 10^8$ MC steps was performed for each system. The temperature was incremented from 400 K to 510 K in steps of 10 K for $(RE)_{25}$, and 460 K to 570 K with an interval of 10 K for $(RD)_{25}$. The first 20% of the trajectory was used as equilibration and discarded from the rest of the analysis.

### 2. Setup of umbrella sampling

Umbrella sampling was used to obtain the free energy profiles as a function of $R_g$. A harmonic biasing potential $U_{bias} = k\left(R_g - R_g^{ref,i}\right)^2$ was applied to restrain the conformations of polyampholytic IDPs within each window, where $k$ was set to be 2.0 kcal/(mol·Å$^2$) and $R_g^{ref,i}$ is the reference $R_g$ for the $i^{th}$ window. The spacing between adjacent windows was 1 Å. The window centers are listed in Table S3. Within each window, MC simulations were performed to sample the conformations. For all the systems except $(GKESKE)_{10}$ , thermal replica exchange Monte Carlo (REMC) simulations were used to enhance the sampling of globular structures in the windows with $R_g^{ref,i} \leq 16$Å. Multiple independent REMC simulations, ranging from 6 to 24 simulations, were performed for each window. Each independent REMC simulation consisted of $4 \times 10^7$MC steps for each temperature. The temperature schedules for the REMC simulations are listed in Table S4. When $R_g^{ref,i} > 16$ Å, standard Metropolis Monte Carlo simulations were performed at certain temperatures. For each window, multiple independent simulations, ranging from 2 to 20 simulations, were performed at each temperature and each independent simulation consisted of $6 \times 10^7$MC steps. The distributions of $R_g$ at each window at different temperatures for all the systems are shown from Figs. S19 to S29. The bin width for the population distribution was set as 0.1 Å. For $(GKESKE)_{10}$, REMC simulations were performed in the windows with $R_g^{ref,i} \leq 20$Å.

The weighted histogram analysis method (WHAM) (7, 8) was used to derive the free energy profile $W(R_g)$. Then, Monte Carlo Bootstrap Error Analysis suggested by Grossfield (9) was used to estimate the statistical uncertainty of the free energy profile. For the $i^{th}$ simulation window, we obtained the $R_g$ distribution from umbrella sampling $P_i^{ub}(R_g)$. Based on this probability distribution, we performed $10^3$ steps of Metropolis Monte Carlo moves and generated a new histogram $p_i^{boot}(R_g)$. Then, we performed WHAM on these new histograms to obtain the free energy profile $W^{boot}(R_g)$. By iterating this process 10 times, we obtained 10 different profiles for $W^{boot}(R_g)$. The standard deviation of the $W^{boot}(R_g)$ profiles was used as the statistical uncertainty of the free energy profile $W(R_g)$. Fig. S5 shows the statistical uncertainty for $(GKESKE)_7$ at 320 K. Table S5 lists the mean statistical uncertainty $\overline{\Delta W}$ for all simulated systems. $\overline{\Delta W}$ is defined as the average uncertainty for all the bins, $\overline{\Delta W} = \sum_{m=1}^{M} \frac{\Delta W_m}{N}$, where $M$ is the total number of bins, and $\Delta W_m$ is the statistical uncertainty of free energy in the $m^{th}$ bin.

### 3. The density peak clustering

We used the density peak clustering algorithm (10, 11) on the probability distribution $P(x)$ to test the hypothesis that distributions obtained from simulations for polyampholytic IDPs comprise two distinct stable states. Cluster centers are characterized by a higher density than their neighbors and by a relatively large distance from points with higher densities. The algorithm groups the samples into different clusters based on two quantities: 1) the local density $\rho_i$ of sample $i$. 2) the minimum distance between the sample $i$ and any sample with higher density $\delta_i = \min_{\rho_j > \rho_i}(d_{ij})$. For the sample with the highest density, we have $\delta_i = \max(d_{ij})$. The decision graph, namely, the plots of $\rho$ against $\delta$, was used to determine the cluster centers and cluster number. The samples with higher $\rho$ and larger $\delta$ than manually chosen cutoff values were chosen as the cluster centers.

In our case, we used the bins of the probability distribution $P(x)$ as the samples for the clustering. The distance between two bins was calculated by taking $d_{ij} = |x_i - x_j|$, where $x_i$ and $x_j$ are the centers of bin $i$ and bin $j$, respectively. For bin $i$, $\rho_i = P_i(x)$, where $P_i(x)$ is the probability of bin $i$. The bin width equals $\frac{0.01}{N^{0.5}}$ Å where $N$ is the number of residues in the peptide and the summation of $P_i(x)$ equals 1.

Fig. S3 and S4 show the decision graphs for $(GKESKE)_7$ and $(GkeSke)_7$, respectively. The decision graph for $(GKESKE)_7$ at 320 K clearly shows two points with high $\rho$ and large $\delta$, and this is indicative of a distribution with two distinct peaks. However, the decision graphs show only one point with both with high $\rho$ and large $\delta$ for all the other probability distributions, indicating one cluster.

### 4. One dimensional Langevin dynamics simulation and mean first passage time calculation

To study the dynamics of state-to-state transitions, we performed a series of one-dimensional Langevin dynamics simulations based on the free energy profile obtained as a function of $R_g$. The process of the diffusion of a single point particle on the one-dimensional free energy profile was simulated. The integration algorithm proposed by Berendsen et al. (12) was used. We used reduced units for the simulations. The free energy from umbrella sampling was divided by $kT_{ref}$ ($T_{ref} = 298\ K$) and the distance was divided by 1 Å, so that 1.0 in the Langevin dynamics simulations corresponds to a length scale of 1 Å. Then, the resulting free energy profile was used as the potential energy profile for the Langevin dynamics simulation. The mass of the particle was set to 1.0, $kT$ was set as 1.0, and the integration time step was 0.05. We performed 17 independent simulations with different initial positions ranging from 10 to 32 and in steps of 2.0. Each independent simulation featured $2 \times 10^8$ steps.

We used the method of Rosta et al., (13) to calculate the first passage time distributions and mean first passage times from the simulation trajectories. These quantify the transitions from globules to coils and vice versa. For a given trajectory, each frame was mapped to a specific state based on our partitioning of the phase space. For a trajectory, the mean first passage time from state A to B is the time taken for a process starting from state A to arrive at state B. To illustrate this, we consider the following short trajectory:

$$1(\mathrm{O}) \to 2(\mathrm{O}) \to 3(\mathrm{A}) \to 4(\mathrm{A}) \to 5(\mathrm{A}) \to 6(\mathrm{O}) \to 7(\mathrm{B}) \to 8(\mathrm{B}) \to 9(\mathrm{A}) \to 10(\mathrm{A}) \to 11(\mathrm{O}) \to 12(\mathrm{B}) \to 13(\mathrm{A}) \to 14(\mathrm{B})$$

where the number indicates the time step, the letter in the bracket indicates the state the process is in at that time. A, B and O represent state A, state B and other states, respectively. The first passage time (FPT) from state A to state B for this short trajectory then includes the following:

- $3(\mathrm{A}) \to 7(\mathrm{B}), \mathrm{FPT} = 4$
- $4(\mathrm{A}) \to 7(\mathrm{B}), \mathrm{FPT} = 3$
- $5(\mathrm{A}) \to 7(\mathrm{B}), \mathrm{FPT} = 2$
- $9(\mathrm{A}) \to 12(\mathrm{B}), \mathrm{FPT} = 3$
- $10(\mathrm{A}) \to 12(\mathrm{B}), \mathrm{FPT} = 2$
- $13(\mathrm{A}) \to 14(\mathrm{B}), \mathrm{FPT} = 1$

The probability of realizing a first passage time that is longer than a lag time *t* can be fit to an exponential equation of the form: $p(FPT > t) = \exp{(-t/\tau)}$ (14). Here, τ is the characteristic time scale for the transition in question.

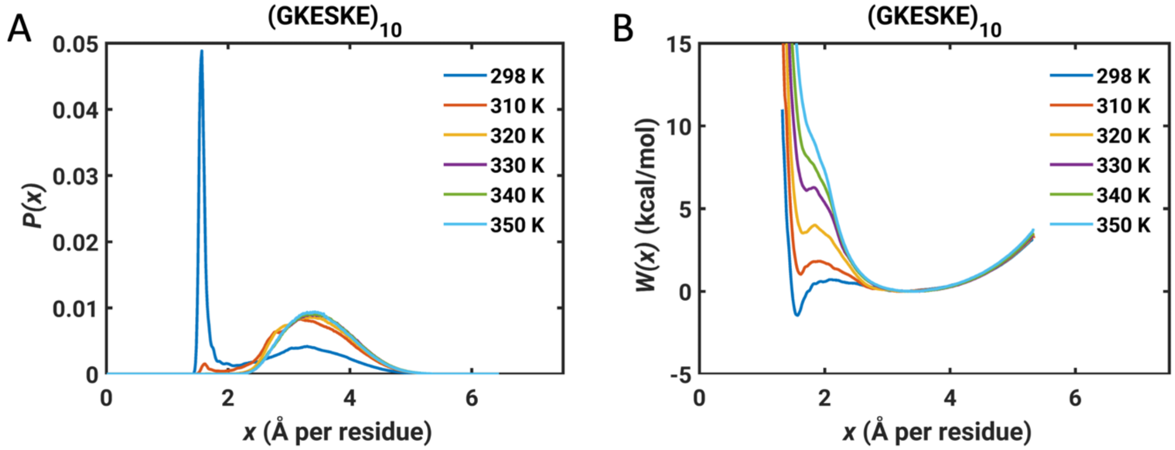


**Fig. S1.** (A) Probability distribution $P(x)$ and (B) free energy profile $W(x)$ as a function of $x$ for $(GKESKE)_{10}$. The ensemble-averaged $x$ at 298 K is 2.48 Å.

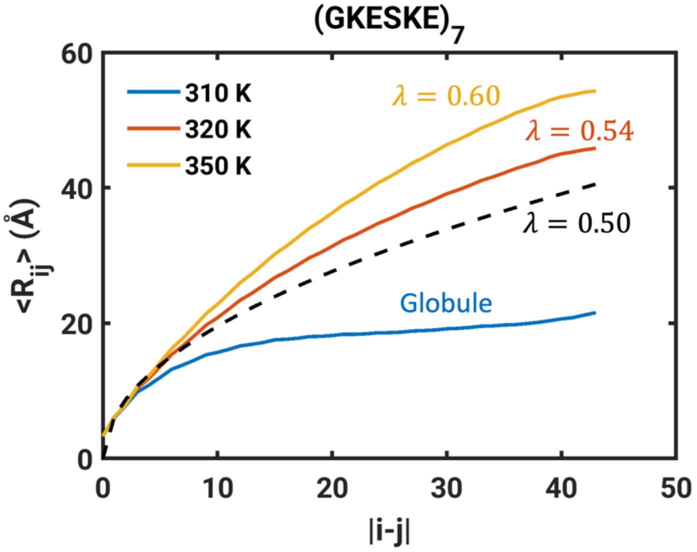


**Fig. S2. The scaling of ensemble-averaged internal distances $\langle R_{ij} \rangle$ with sequence separation |j-i| for (GKESKE)$_7$ at different temperatures.** The black dashed line shows the reference internal scaling distance corresponding to λ=0.5, namely $R_{ij} = R_0|i-j|^{0.5}$. The pre-factor $R_0$ is the value corresponding to |*i*-*j*|=1, which is obtained from simulations. This shows very weak temperature dependence. The scaling exponent is 0.54 at 320 K. At the high temperature of 350K, the peptide behaves like a self-avoiding walk with a scaling exponent of 0.60. The scaling exponent is fitted using the data when |*i*-*j*|≥10. The internal distance is obtained by reweighting the biased simulations $< R_{ij} > = \frac{\sum_{n=1}^{N}\left[p(x_n)\times\sum_{m=1}^{M}\frac{R_{ij}(x_n,m)}{M}\right]}{\sum_{n=1}^{N}p(x_n)}$, where $p(x_n)$ is the population for the bin with center of $x_n$ from WHAM. The bin width is 0.1 Å. M random conformations are chosen for each bin. $R_{ij}(x_n, m)$ is the internal distance for the *m*th conformation in the bin with center of $x_n$. M is set to 100.At the lower temperature of 310 K, equilibrium globules form, and this is associated with a plateauing behavior of the internal scaling profiles (15). Contrary to recent approaches prescribed in the literature (16), the internal scaling profiles for globules cannot be modeled as fractals and one should not extract scaling exponents from such profiles (15) because this corresponds to the breaking of dilatation symmetry. We assign simulation temperatures corresponding to the plateauing regime of the internal scaling profiles as being temperatures corresponding to pure globules. Having assigned a temperature corresponding to the globule regime, we identify the first peak for the $R_g$ distribution at low temperatures as the stable globule, and this corresponds to $x$ = 1.6 Å. This is shown in Fig. 1B. For the higher temperatures, we use the internal scaling profile to extract the scaling exponent and find that this corresponds to that of SAWs. We associate the second peak located at $x$=3.4 Å as the second stable state corresponding to SAWs.

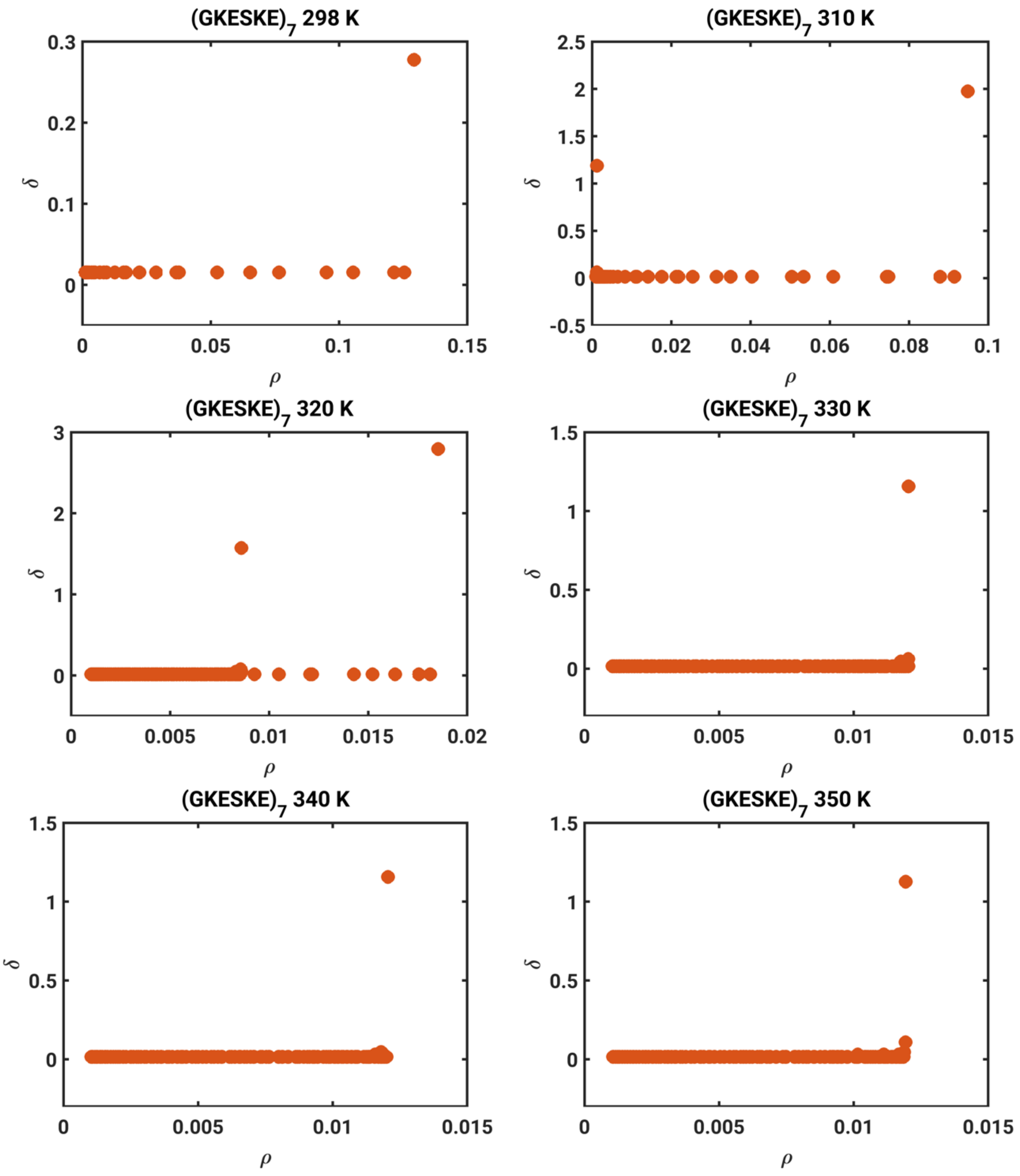


**Fig. S3. The decision graphs of density peak clustering applied to analyze the probability distribution *P*(*x*) for (GKESKE)$_7$ at different temperatures.** $P(x)$ at 320 K shows two clusters with large $\rho$ and $\delta$.

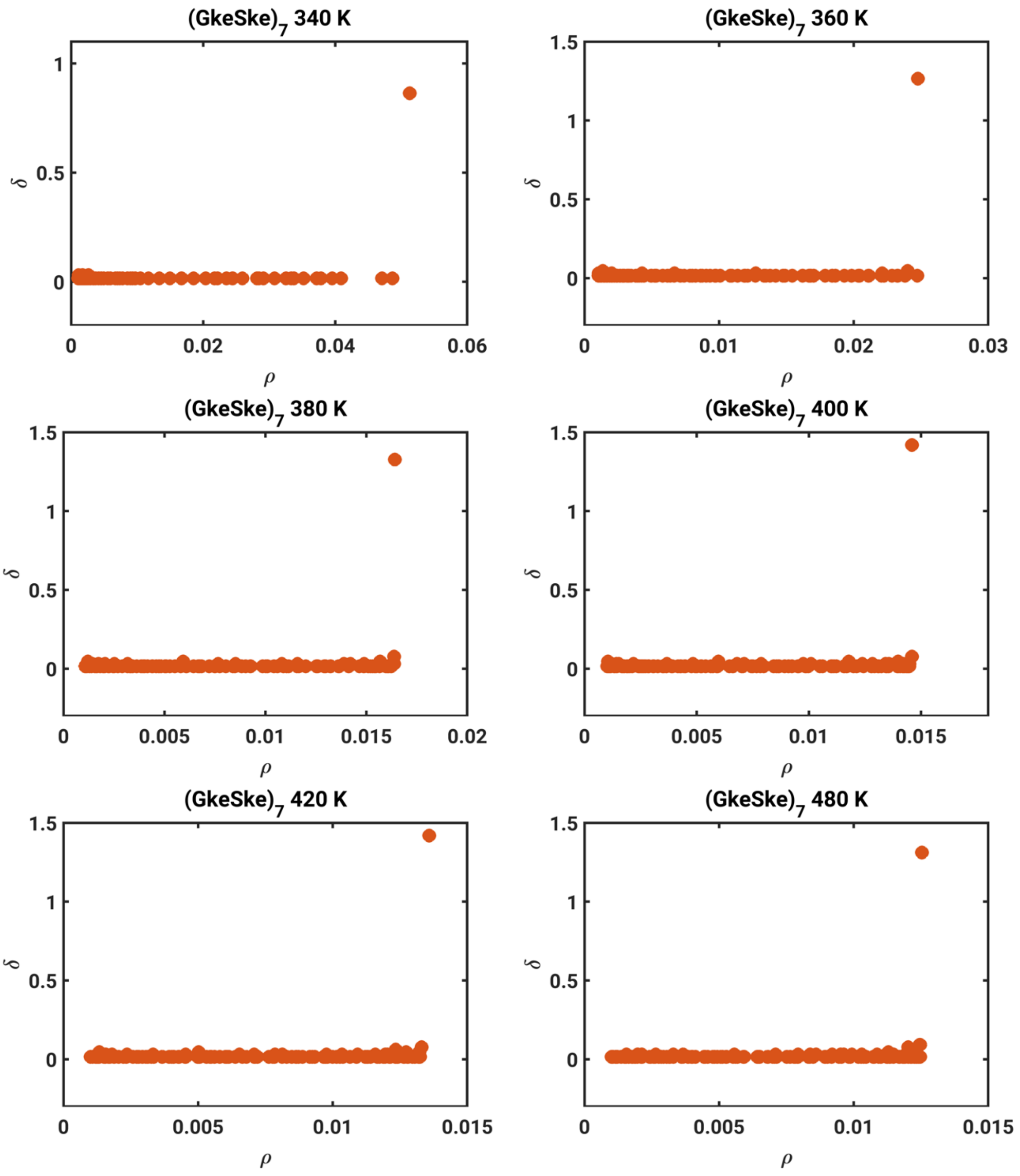


**Fig. S4. The decision graphs of density peak clustering applied to analyze the probability distribution *P*(*x*) for (GkeSke)$_7$ at different temperatures.** *P*(*x*) at all these temperatures shows one cluster.

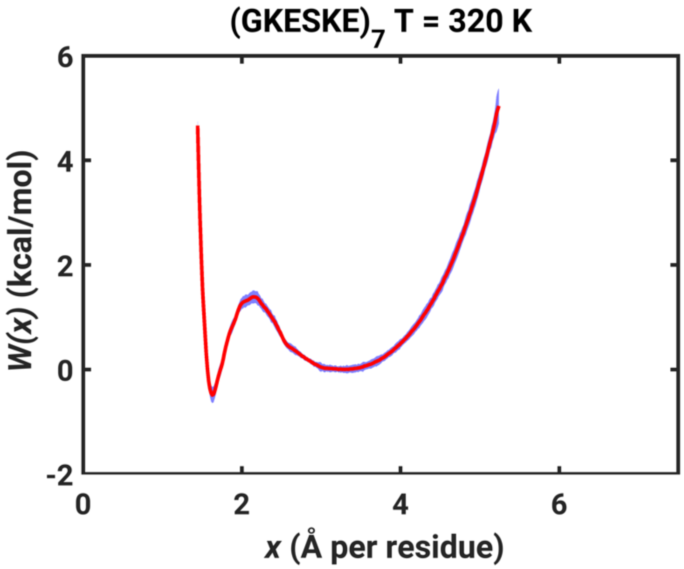


**Fig. S5. The free energy profile *W*(*x*) for $(GKESKE)_7$ at 320 K (in red) along with the uncertainty in *W*(*x*) (in blue) as estimated from bootstrapping.**

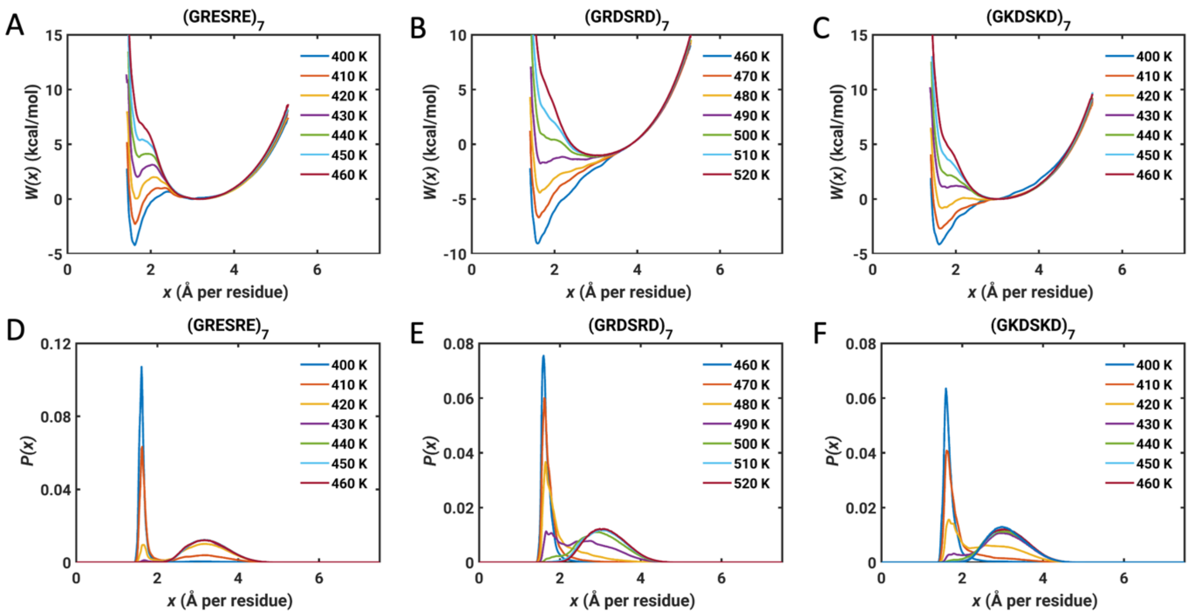


**Fig. S6. The free energy profiles *W*(*x*) and probability distributions *P*(*x*) as a function of *x*, for (GRESRE)$_7$, (GRDSRD)$_7$ and (GKDSKD)$_7$.**

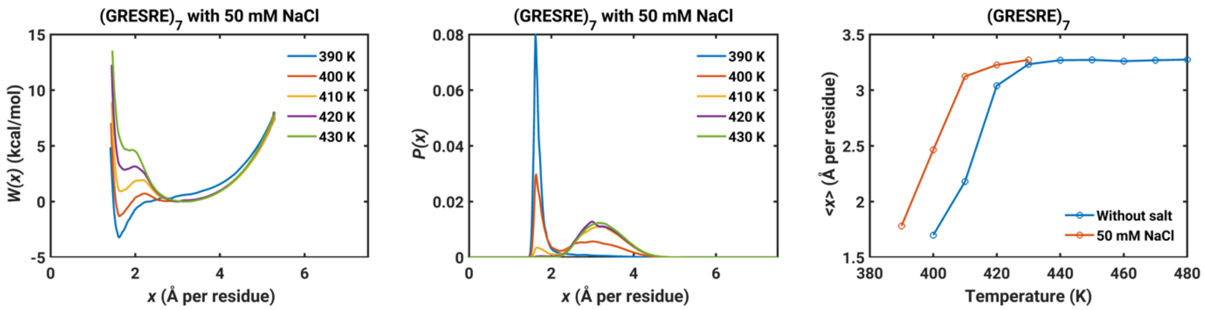


**Fig. S7. Increasing the concentration of NaCl, as an exemplar of monovalent salts, decreases the temperature of bistability.** The free energy profile $W(x)$ (left panel) and probability distribution $P(x)$ (middle panel) for $(GRESRE)_7$ with 50 mM NaCl. Right panel shows the normalized $R_g$ as a function of temperature for $(GRESRE)_7$ without salt and in the presence of 50 mM NaCl. Parameters for the monovalent ions were based on those of Mao and Pappu (17).

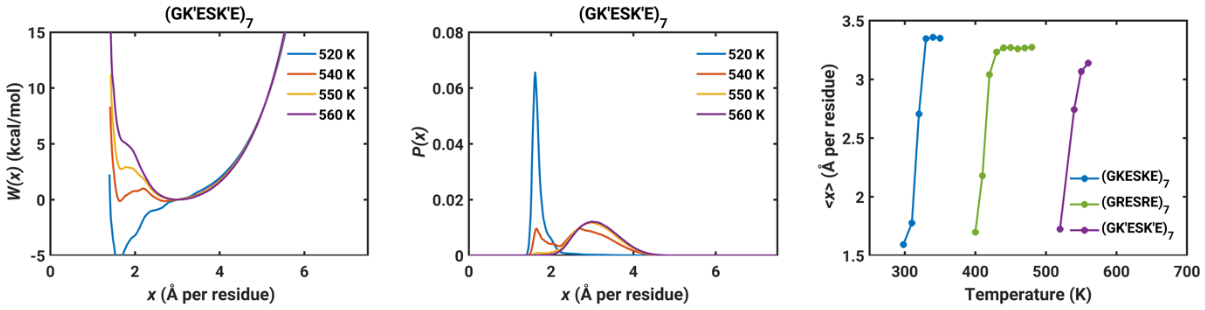


**Fig. S8. The free energy profile *W*(*x*) (left panel) and probability distribution *P*(*x*) (middle panel) for (GK'ESK'E)$_7$.** Here, K' denotes Lys with the free energy of hydration of Arg. Right panel shows the normalized $R_g$ as a function of temperature for (GKESKE)$_7$, (GRESRE)$_7$ and (GK'ESK'E)$_7$. These data show that differences in steric volumes contribute to differences between Arg and Lys containing sequences.

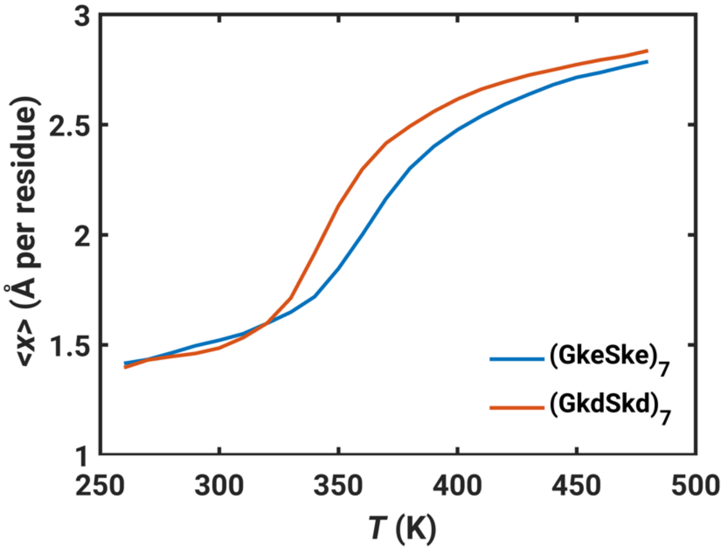


**Fig. S9. Ensemble-averaged values of *x* for sequences devoid of charges.** The data are plotted against simulation temperature for $(GkeSke)_7$ and $(GkdSkd)_7$.

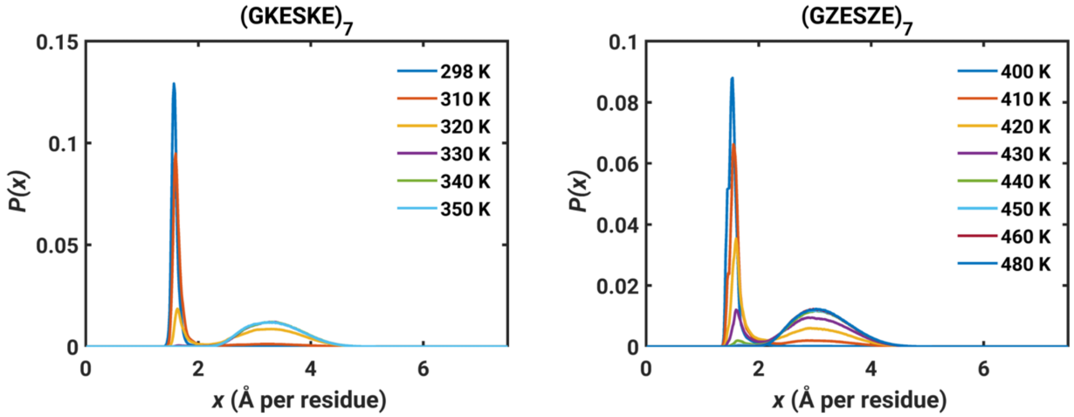


**Fig. S10. Probability distribution *P*(*x*) as a function of *x* for (GKESKE)$_7$ (left) and** (GZESZE)$_7$ **(right).** Here, Z refers to DAB. The transition temperature for (GKESKE)$_7$ and (GZESZE)$_7$ are ~315 K and ~420 K, respectively.

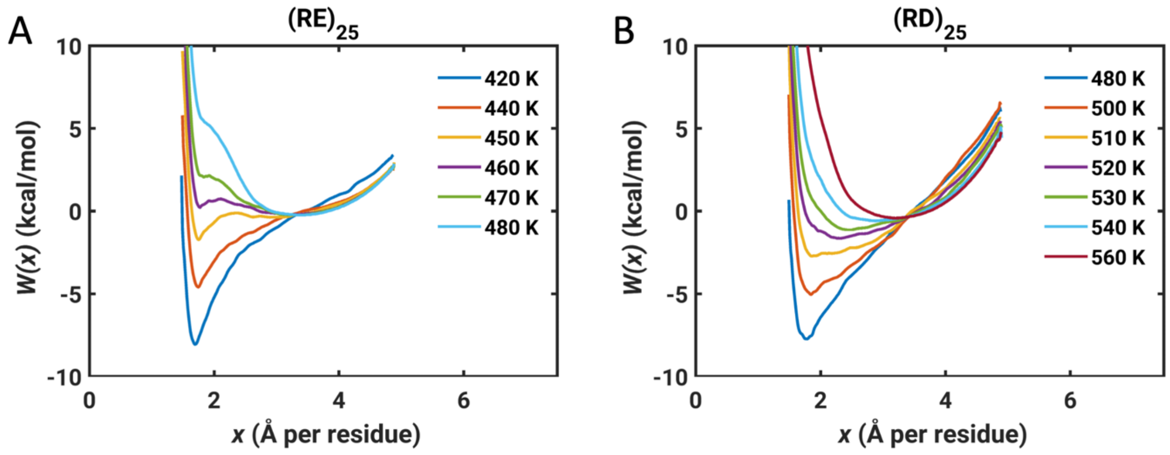


**Fig. S11. Free energy profile *W*(*x*) as a function of *x* for (A) $(RE)_{25}$ and (B) $(RD)_{25}$.**

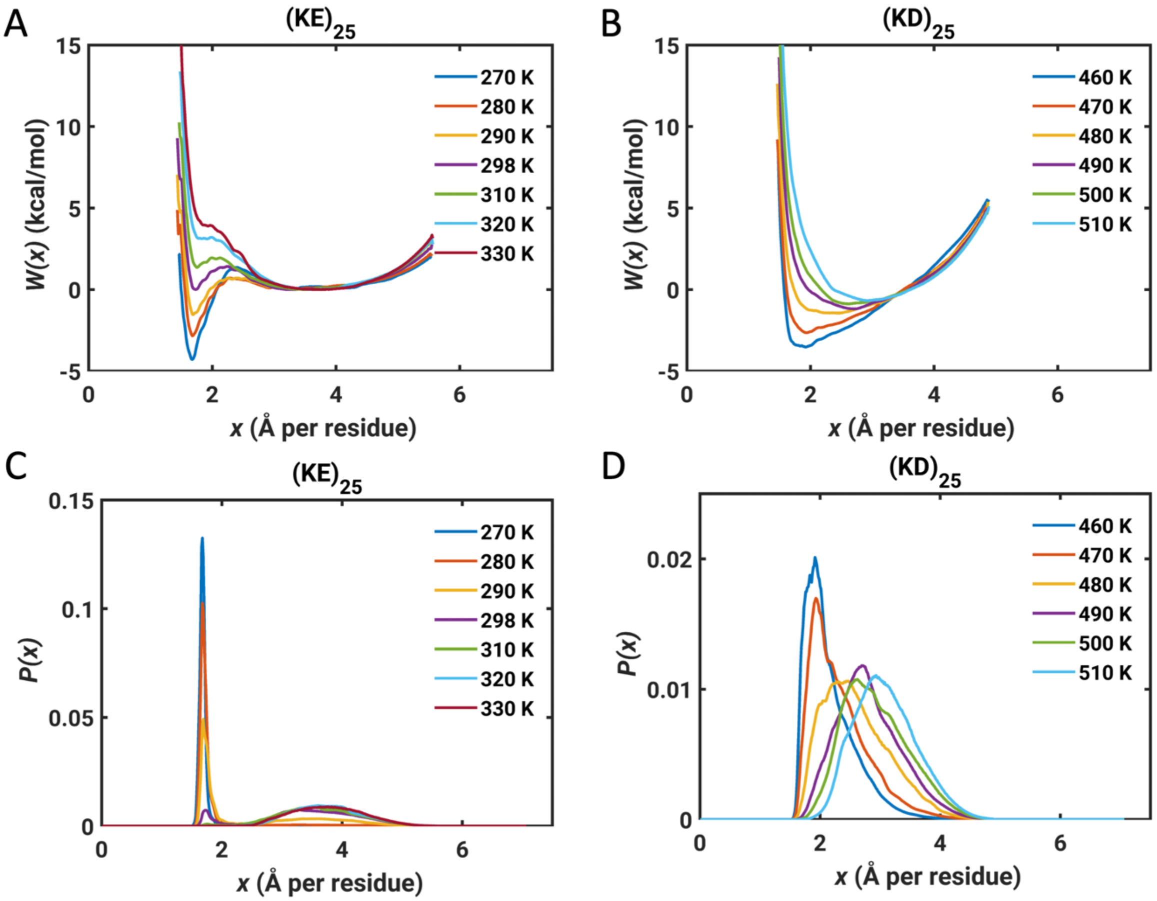


**Fig. S12.** **Free energy profile** ***W*****(*****x*****) and probability distribution** ***P*****(*****x*****) as a function of** ***x*** **for (KE)$_{25}$ (A and C) and (KD)$_{25}$ (B and D).**

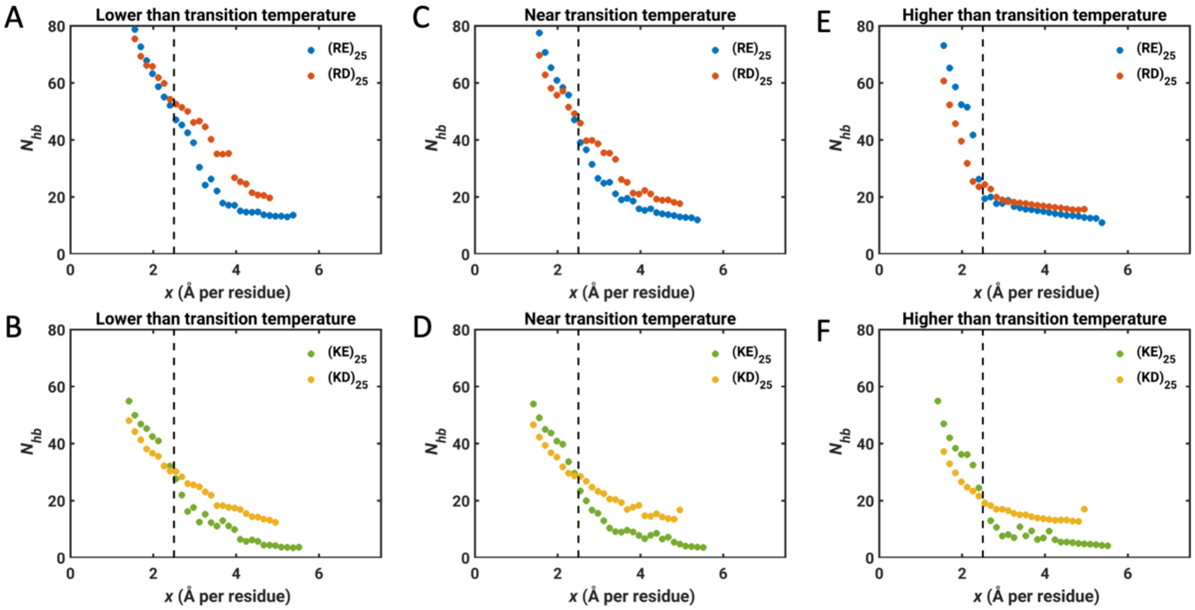


**Fig. S13. Number of hydrogen bonds $N_{hb}$ as a function of $x$, for $(RE)_{25}$, $(RD)_{25}$, $(KE)_{25}$ and $(KD)_{25}$ at the temperature (A-B) lower than transition temperature, (C-D) near the transition temperature and (E-F) higher than the transition temperature.** The three temperatures are 440 K, 450 K and 480 K for $(RE)_{25}$, 480 K, 510 K and 560 K for $(RD)_{25}$, 280 K, 290 K and 320 K for $(KE)_{25}$, 460 K, 470 K and 510 K for $(KD)_{25}$. The **baker_hubbard** function implemented in MDTraj (18) was used to identify the hydrogen bonds. The default criterion for a hydrogen bond was used; the angle between the donor-H acceptor is larger than 120° and that the distance between H and acceptor is less than 2.5 Å. Umbrella sampling trajectories were used to calculate $N_{hb}$.

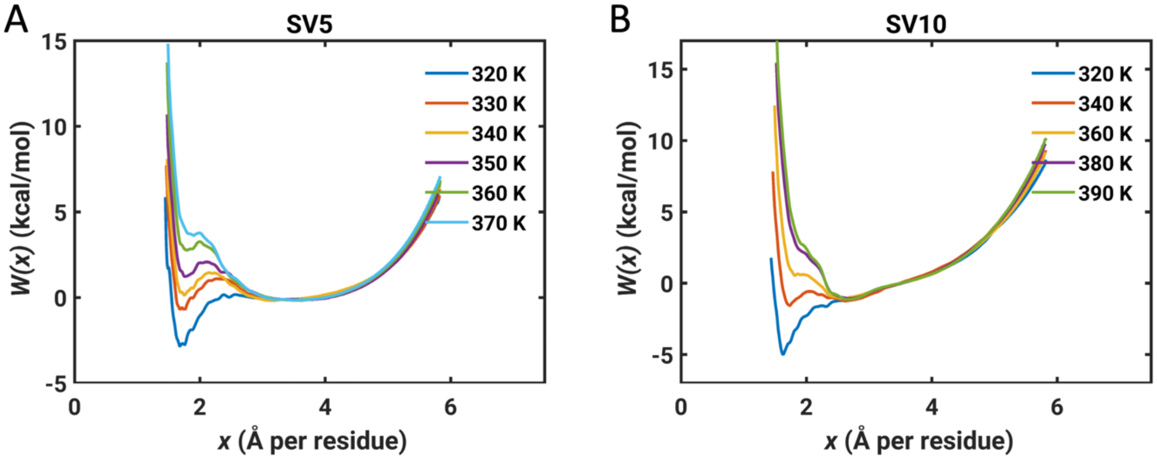


**Fig. S14. Free energy profile *W*(*x*) as a function of *x* for (A) sv5 and (B) sv10.**

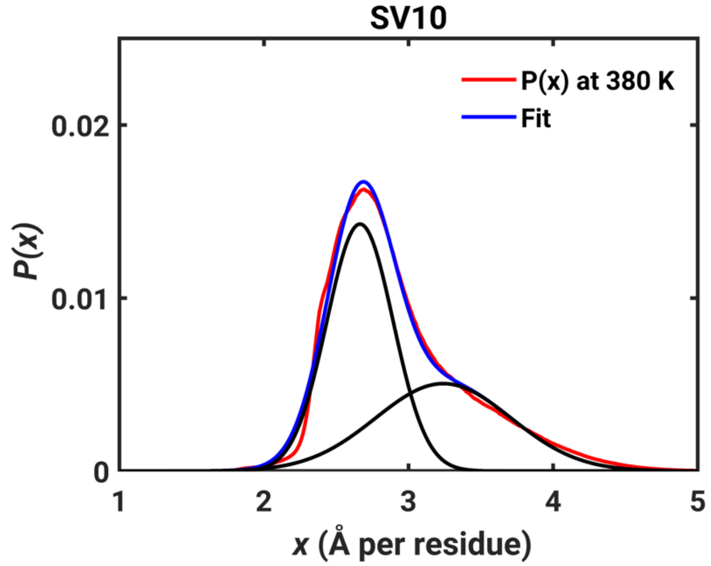


**Fig. S15. The probability distribution *P*(*x*) at 380 K for sv10 (red) was fit to a sum of two Gaussian distributions (black).** The summation of the two Gaussian distributions is shown in the blue profile: $P(x) = 0.1008 \times \exp\left[-\left(\frac{x-2.6644}{0.3240}\right)^2\right] + 0.0357 \times \exp\left[-\left(\frac{x-3.2385}{0.6657}\right)^2\right]$. The relative populations in the Gaussian distribution with the center at 2.6644 Å and 3.2385 Å are 0.5785 and 0.4214, respectively.

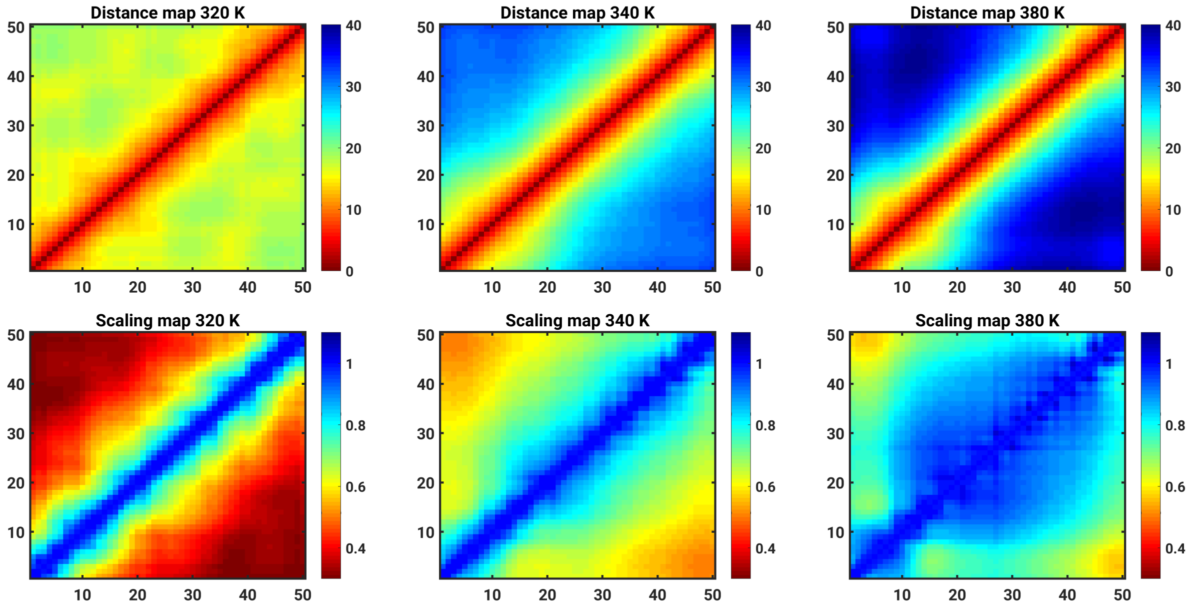


**Fig. S16. Distance maps (upper panel) and scaling maps (lower panel) for SV10 at 320 K, 340 K and 380 K.** These three temperatures favor the globule state, the bistable state, and the coil state, respectively. The distance map represents the averaged distance between each pair of residues. The scaling map is obtained by normalizing the distance map using the inter-residue distances from a simulation performed in the excluded volume limit (15) for SV10. The distance map $D_{rwt}$ is obtained by reweighting the biased simulation data: $D_{rwt} = \frac{\sum_{i=1}^{N}\left[p(x_i)\times\sum_{j=1}^{M}\frac{D(x_i,j)}{M}\right]}{\sum_{i=1}^{N}p(x_i)}$, where $p(x_i)$ is the population for the bin with center of $x_i$ from WHAM. The bin width is 0.1 Å. M random conformations are chosen for each bin. $D(x_i, j)$ is the distance map for the $j^{\text{th}}$ conformation in the bin with center of $x_i$. M is set to 100.

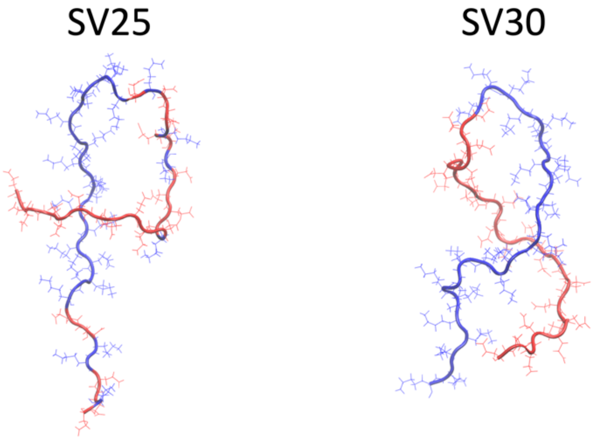


**Fig. S17. Examples of structures stabilized by electrostatic interactions for sequence sv25 and sv30.** Nomenclature as in Das and Pappu (19). Lys and Asp residues are colored by blue and red, respectively. VMD (20) was used for visualization.

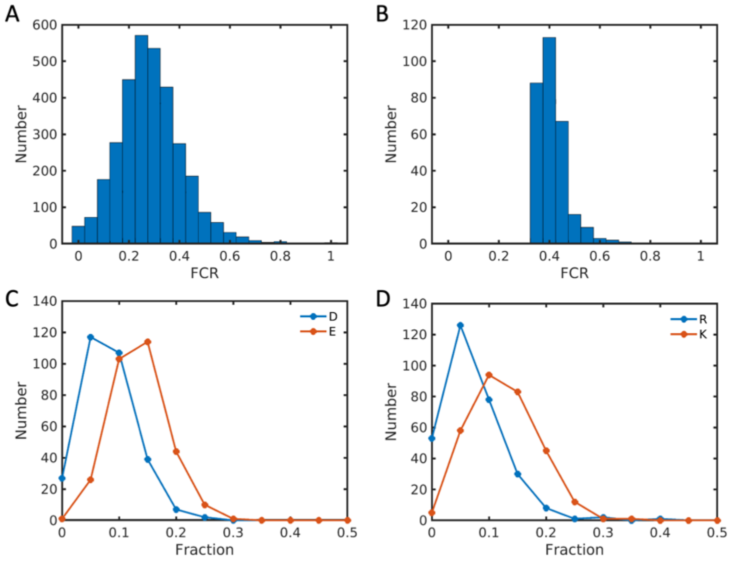


**Fig. S18. Distribution of fractions of charged residues (FCR) for (A) IDRs with length ≥ 30 in DisProt 2022-03 database (21-23) and (B) strong polyampholytes with length ≥ 30.** In the DisProt database, one protein may have multiple IDRs termed as fragments. If one fragment is a subset of another one, we only choose the longer one for the analysis. Strong polyampholytes are sequences with NCPR ≤ |0.2| and FCR > 0.35. A total of 202 out of 3236 IDRs with length ≥30 have FCR equal or larger than 0.5. 26 out of 299 strong polyampholytes with length ≥30 have FCR values equal to or larger than 0.5. Distribution of fraction of (C) Asp or Glu, (D) Arg or Lys in the strong polyampholytes.

**Table S1: Summary of inputs to TCPD approach used for estimating values free energies of hydration for model compounds that mimic charged versions of $Arg^+$, $Lys^+$, $Asp^-$, and $Glu^-$.** The values different from those prescribed by Fossat et al. (4) are highlighted in blue. $-252.7$ kcal/mol is used for the proton free energy of hydration ($\Delta u_h^{H^+}$).

| **Sidechain mimicked by the model compound** | **$pK_a$** | **Gas phase basicity (kcal / mol)** | **$\Delta\mu_h^B, \Delta\mu_h^{AH}$ (kcal / mol)** |
|---|---|---|---|
| $Arg^+$ | 13.9 | 234.6 | -10.0 |
| $Lys^+$ | 10.7 | 211.3 | -4.3 |
| $Asp^-$ | 4.76 | 341.4 | -6.7 |
| $Glu^-$ | 4.88 | 340.4 | -6.5 |

**Table S2. Reference free energies of hydration (ΔG°) for the sidechains of amino acids used in this work.**

| | **$Arg^+$** | **$Lys^+$** | **$Asp^-$** | **$Glu^-$** | **Dab** | **Neutral Lys (k)** | **Neutral Asp (d)** | **Neutral Glu (e)** |
|---|---|---|---|---|---|---|---|---|
| ΔG° (kcal/mol) | -58.54 | -71.82 | -100.40 | -99.04 | -71.82 | -4.3 | -6.7 | -6.5 |

**Table S3. Number of windows for the umbrella sampling and the smallest and largest window centers for each system.** The spacing between pairs of adjacent windows was 1 Å.

| System | N | $R_g^{ref,1}$ (Å) | $R_g^{ref,N}$ (Å) |
|---|---|---|---|
| $(GKESKE)_{10}$ | 31 | 10 | 40 |
| $(GKESKE)_7$ | 25 | 9 | 33 |
| $(GKDSKD)_7$ | 25 | 9 | 33 |
| $(GRESRE)_7$ | 25 | 9 | 33 |
| $(GRDSRD)_7$ | 25 | 9 | 33 |
| $(KE)_{25}$ | 29 | 10 | 38 |
| $(KD)_{25}$ | 24 | 10 | 33 |
| $(RE)_{25}$ | 24 | 10 | 33 |
| $(RD)_{25}$ | 24 | 10 | 33 |
| SV5 | 31 | 10 | 40 |
| SV10 | 31 | 10 | 40 |

**Table S4. Temperature schedule for the REMC simulation in windows with $R_g^{ref,i} \leq 16$ Å.** Note that the temperature schedule for $(GKESKE)_{10}$ is the same as for $(GKESKE)_7$. REMC simulations were performed in windows with $R_g^{ref,i} \leq 20$ Å for $(GKESKE)_{10}$.

| System | Temperatures (K) |
|---|---|
| $(GKESKE)_7$ | 298, 310, 320, 330, 340, 350 |
| $(GKDSKD)_7$ | 400 to 470 with an interval of 10 |
| $(GRESRE)_7$ | 400 to 480 with an interval of 10 |
| $(GRDSRD)_7$ | 460 to 530 with an interval of 10 |
| $(KE)_{25}$ | 270, 280, 290, 298, 310, 320, 330, 340, 350 |
| $(KD)_{25}$ | 460 to 510 with an interval of 10 |
| $(RE)_{25}$ | 420 to 490 with an interval of 10 |
| $(RD)_{25}$ | 470 to 560 with an interval of 10 |
| SV5 | 320 to 390 with an interval of 10 |
| SV10 | 320 to 390 with an interval of 10 |

**Table S5. The mean uncertainty in the free energy profile $\overline{\Delta W}$ at different temperatures for the simulated systems.** The mean uncertainty was computed as the average value of uncertainties at different positions, which are estimated from Monte-Carlo bootstrapping (details above).

| | | | | | | | | |
|---|---|---|---|---|---|---|---|---|
| **$(GKESKE)_7$** | T (K) | 298 | 310 | 320 | 330 | 340 | 350 | |
| | $\overline{\Delta W}$ (kcal/mol) | 0.11 | 0.16 | 0.11 | 0.11 | 0.12 | 0.13 | |
| **$(GKDSKD)_7$** | T (K) | 400 | 410 | 420 | 430 | 440 | 450 | 460 |
| | $\overline{\Delta W}$ (kcal/mol) | 0.17 | 0.14 | 0.14 | 0.14 | 0.13 | 0.14 | 0.15 |
| **$(GRESRE)_7$** | T (K) | 400 | 410 | 420 | 430 | 440 | 450 | 460 |
| | $\overline{\Delta W}$ (kcal/mol) | 0.16 | 0.16 | 0.12 | 0.13 | 0.12 | 0.15 | 0.16 |
| **$(GRDSRD)_7$** | T (K) | 460 | 470 | 480 | 490 | 500 | 510 | 520 |
| | $\overline{\Delta W}$ (kcal/mol) | 0.28 | 0.19 | 0.19 | 0.20 | 0.15 | 0.15 | 0.18 |
| **$(KE)_{25}$** | T (K) | 270 | 280 | 290 | 298 | 310 | 320 | 330 |
| | $\overline{\Delta W}$ (kcal/mol) | 0.14 | 0.18 | 0.17 | 0.10 | 0.09 | 0.12 | 0.10 |
| **$(KD)_{25}$** | T (K) | 460 | 470 | 480 | 490 | 500 | 510 | |
| | $\overline{\Delta W}$ (kcal/mol) | 0.14 | 0.14 | 0.16 | 0.14 | 0.16 | 0.14 | |
| **$(RE)_{25}$** | T (K) | 420 | 440 | 450 | 460 | 470 | 480 | |
| | $\overline{\Delta W}$ (kcal/mol) | 0.20 | 0.19 | 0.17 | 0.11 | 0.13 | 0.15 | |
| **$(RD)_{25}$** | T (K) | 480 | 500 | 510 | 520 | 530 | 540 | 560 |
| | $\overline{\Delta W}$ (kcal/mol) | 0.24 | 0.22 | 0.22 | 0.19 | 0.15 | 0.19 | 0.15 |
| **SV5** | T (K) | 320 | 330 | 340 | 350 | 360 | 370 | |
| | $\overline{\Delta W}$ (kcal/mol) | 0.20 | 0.13 | 0.10 | 0.13 | 0.17 | 0.14 | |
| **SV10** | T (K) | 320 | 340 | 360 | 380 | 390 | | |
| | $\overline{\Delta W}$ (kcal/mol) | 0.21 | 0.11 | 0.12 | 0.16 | 0.15 | | |

**Table S6. Top 15 strong polyampholytes with length ≥ 30 ranked by FCR from DisProt 2022-03 database.** The sequence, FCR and NPCR of the IDR and the protein which the IDR belongs to are listed. The disorder function, molecular function, biological process, or structural transition which the IDR involves in is also listed if available in the database.

| | | | |
|---|---|---|---|
| DisProt ID: **DP02869** | Fragment: 481-551 | FCR: 0.69 | NPCR: 0.15 |
| Protein name | Cleavage and polyadenylation specificity factor subunit 6 | | |
| Sequence | ESKSYGSGSRRERSRERDHSRSREKSRRHKSRSRDRHDDYYRERSRERERH RDRDRDRDRERDREREYRHR | | |
| Disorder function | **Regulation of phosphorylation**, IDPO:00025 (Fragment: 492-496, 498-502, 509-515) | | |
| Biological process | Localization, GO:0051179 (Fragment: 481-551) | | |

| | | | |
|---|---|---|---|
| DisProt ID: **DP00075** | Fragment: 175-214 | FCR: 0.65 | NPCR: 0.10 |
| Protein name | DNA topoisomerase 1 | | |
| Sequence | KPKNKDKDKKVPEPDNKKKKPKKEEEQKWKWWEEERYPEG | | |

| | | | |
|---|---|---|---|
| DisProt ID: **DP02332** | Fragment: 1-257 | FCR: 0.63 | NPCR: 0.10 |
| Protein name | Probable ATP-dependent RNA helicase DDX23 | | |
| Sequence | MAGELADKKDRDASPSKEERKRSRTPDRERDRDRDRKSSPSKDRKRHRSR DRRRGGSRSRSRSRSKSAERERRHKERERDKERDRNKKDRDRDKDGHRR DKDRKRSSLSPGRGKDFKSRKDRDSKKDEEDEHGDKKPKAQPLSLEELLA KKKAEEEAEAKPKFLSKAEREAEALKRRQQEVEERQRMLEEERKKRKQF QDLGRKMLEDPQERERRERRERMERETNGNEDEEGRQKIREEKDKSKELH AIKERYLGG | | |

| | | | |
|---|---|---|---|
| DisProt ID: **DP03459** | Fragment: 381-416 | FCR: 0.61 | NPCR: -0.06 |
| Protein name | Transcription initiation factor TFIID subunit 5 | | |
| Sequence | EIEVPLDDEDEEGENEEGKPKKKKPKKDSIGSKSKK | | |

| | | | |
|---|---|---|---|
| DisProt ID: **DP02856** | Fragment: 73-107 | FCR: 0.60 | NPCR: -0.03 |
| Protein name | Chromobox protein homolog 1 | | |
| Sequence | TAHETDKSEGGKRKADSDSEDKGEESKPKKKKEES | | |
| Disorder function | **Flexible linker/spacer**, IDPO:00502 (Fragment 97-109) | | |
| Molecular function | Molecular adaptor activity, GO:0060090 (Fragment 97-109) ; DNA binding, GO:0003677 (Fragment 97-109); Molecular function regulator, GO:0098772 (Fragment 97-109) | | |

| | | | |
|---|---|---|---|
| DisProt ID: **DP00075** | Fragment: 1-174 | FCR: 0.59 | NPCR: 0.04 |
| Protein name | DNA topoisomerase 1 | | |
| Sequence | MSGDHLHNDSQIEADFRLNDSHKHKDKHKDREHRHKEHKKEKDREKSKH SNSEHKDSEKKHKEKEKTKHKDGSSEKHKDKHKDRDKEKRKEEKVRASG | | |

| | |
|---|---|
| | DAKIKKEKENGFSSPPQIKDEPEDDGYFVPPKEDIKPLKRPRDEDDADYKP KKIKTEDTKKEKKRKLEEEEDGKLK |
| Molecular function | DNA binding, GO:0003677 (Fragment 1-174), DNA binding, bending, GO:0008301 (Fragment 1-174) |

| DisProt ID: **DP03149** | Fragment: 573-639 | FCR: 0.57 | NPCR: 0.03 |
|---|---|---|---|
| Protein name | N-alpha-acetyltransferase 15, NatA auxiliary subunit | | |
| Sequence | LTDENKEHEADTANMSDKELKKLRNKQRRAQKKAQIEEEKKNAEKEKQQ RNQKKKKDDDDEEIGGPK | | |

| DisProt ID: **DP02171** | Fragment: 184-437 | FCR: 0.56 | NPCR: 0.06 |
|---|---|---|---|
| Protein name | U1 small nuclear ribonucleoprotein 70 kDa | | |
| Sequence | VKGWRPRRLGGGLGGTRRGGADVNIRHSGRDDTSRYDERPGPSPLPHRDR DRDRERERRERSRERDKERERRRSRSRDRRRRSRSRDKEERRRSRERSKDK DRDRKRRSSRSRERARRERERKEELRGGGGDMAEPSEAGDAPPDDGPPGE LGPDGPDGPEEKGRDRDRERRRSHRSERERRRDRDRDRDRDREHKRGERG SERGRDEARGGGGGQDNGLEGLGNDSRDMYMESEGGDGYLAPENGYLM EAAPE | | |
| Molecular function | Protein binding, GO:0005515 (Fragment: 184-437) | | |

| DisProt ID: **DP00217** | Fragment: 120-200 | FCR: 0.56 | NPCR: -0.11 |
|---|---|---|---|
| Protein name | Nucleoplasmin | | |
| Sequence | MEEDYSWAEEEDEGEAEGEEEEEEEEDQESPPKAVKRPAATKKAGQAKK KKLDKEDESSEEDSPTKKGKGAGRGRKPAAKK | | |
| Molecular function | Protein binding, GO:0005515 (Fragment: 120-200, 153-171) | | |

| DisProt ID: **DP02233** | Fragment: 377-482 | FCR: 0.54 | NPCR: -0.14 |
|---|---|---|---|
| Protein name | Histone deacetylase 1 | | |
| Sequence | PGVQMQAIPEDAIPEESGDEDEDDPDKRISICSSDKRIACEEEFSDSEEEGEG GRKNSSNFKKAKRVKTEDEKEKDPEEKKEVTEEEKTKEEKPEAKGVKEEV KLA | | |
| Molecular function | Protein binding, GO:0005515 (Fragment: 377-482) | | |

| DisProt ID: **DP00653** | Fragment: 139-168 | FCR: 0.53 | NPCR: 0.07 |
|---|---|---|---|
| Protein name | Talin-1 | | |
| Sequence | DEGTGTLRKDKTLLRDEKKMEKLKQKLHTD | | |
| Disorder function | **Flexible linker/spacer**, IDPO:00502 (Fragment 139-168) | | |

| DisProt ID: **DP01871** | Fragment: 137-181 | FCR: 0.53 | NPCR: 0.04 |
|---|---|---|---|
| Protein name | Troponin I, fast skeletal muscle | | |
| Sequence | LRANLKQVKKEDTEKEKDLRDVGDWRKNIEEKSGMEGRKKMFEAG | | |

| DisProt ID: **DP02826** | Fragment: 247-280 | FCR: 0.53 | NPCR: -0.18 |
|---|---|---|---|

| | |
|---|---|
| Protein name | Autophagy-related protein 3 |
| Sequence | VRVVRQRRRKELQEEQELDGVGDWEDLQDDIDDS |
| Molecular function | Protein binding, GO:0005515 (Fragment: 269-274) |

| DisProt ID: **DP00307** | Fragment: 288-323 | FCR: 0.53 | NPCR: -0.14 |
|---|---|---|---|
| Protein name | Cyclin-H | | |
| Sequence | NVITKKRKGYEDDDYVSKKSKHEEEEWTDDDLVESL | | |

| DisProt ID: **DP02327** | Fragment: 233-270 | FCR: 0.53 | NPCR: -0.11 |
|---|---|---|---|
| Protein name | Phosphatidylinositol transfer protein alpha isoform | | |
| Sequence | DKWVDLTMDDIRRMEEETKRQLDEMRQKDPVKGMTADD | | |
| Structural transition | Order to pre-molten globule, IDPO:00058 (Fragment: 233-270) | | |

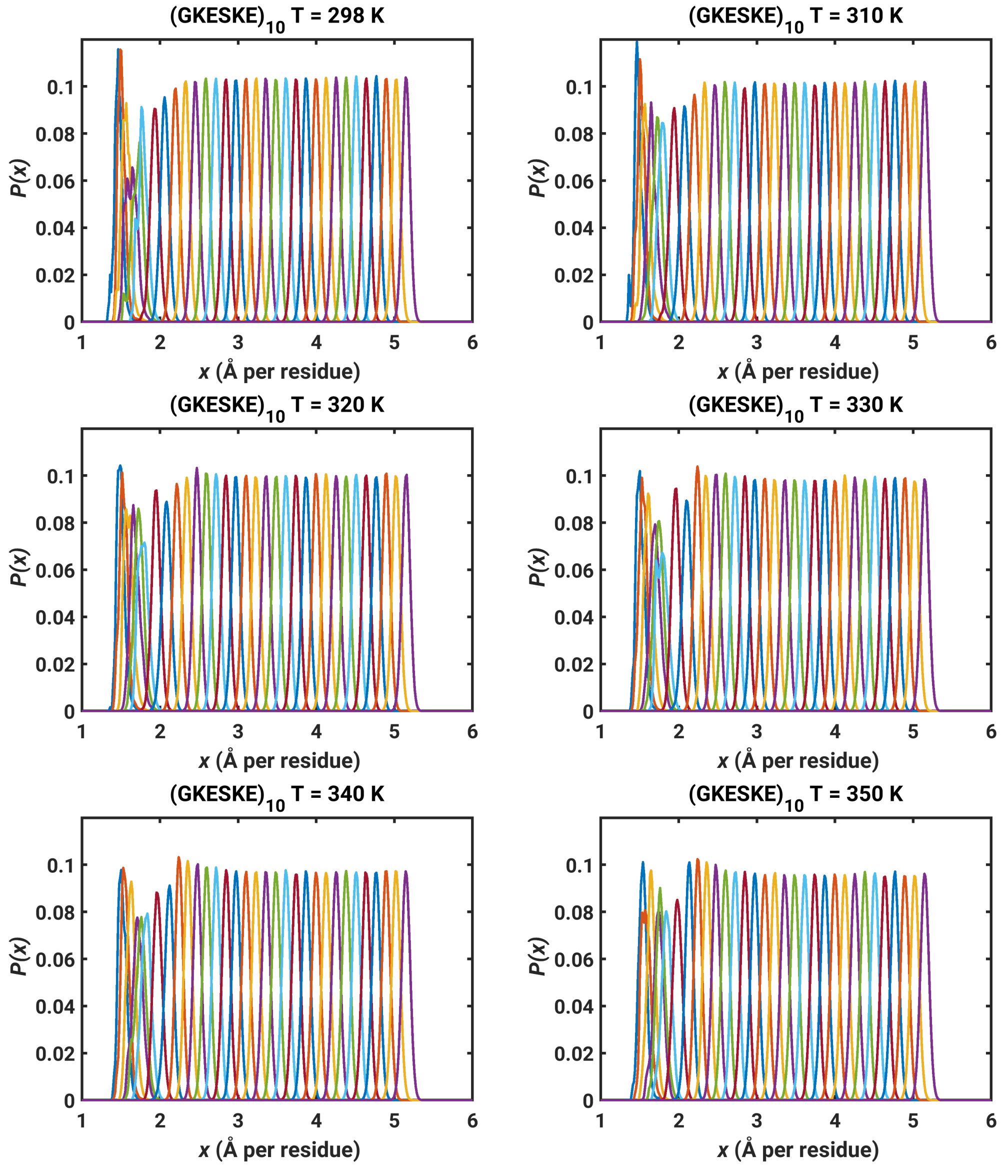


**Fig. S19. Probability distribution *P*(*x*) of *x* in each window of the umbrella sampling for $(GKESKE)_{10}$.**

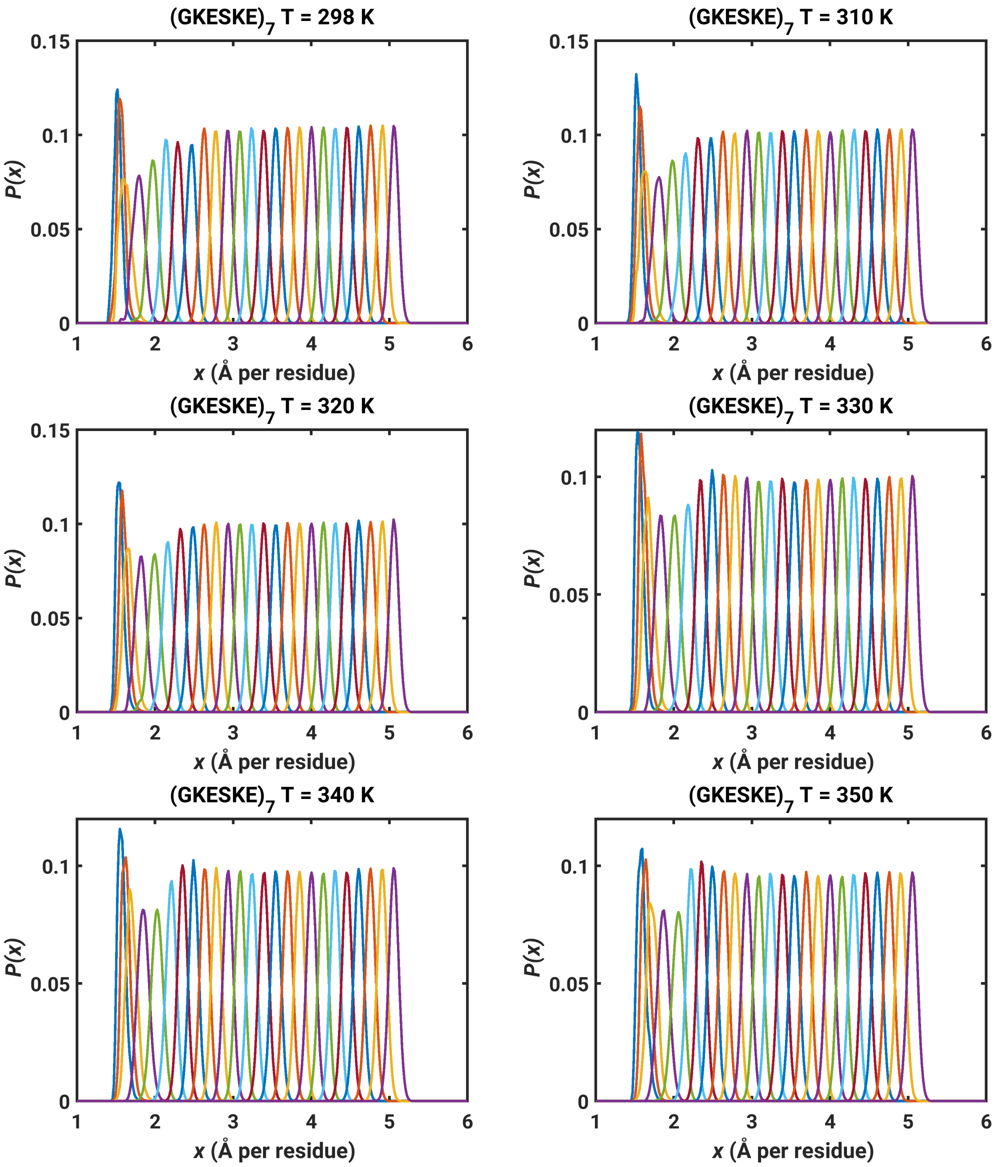


**Fig. S20. Probability distribution $P(x)$ of $x$ in each window of the umbrella sampling for $(GKESKE)_7$.**

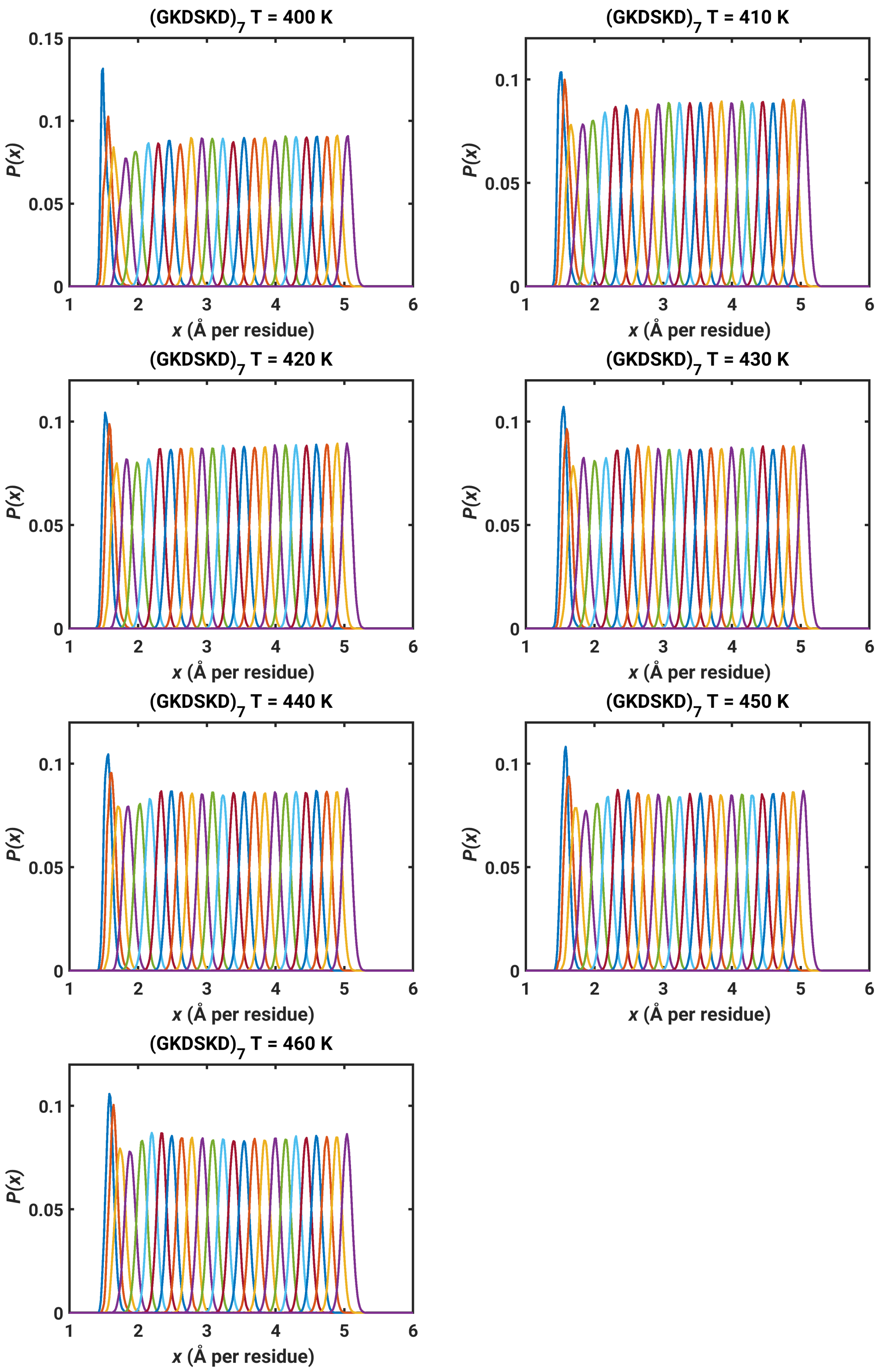


**Fig. S21. Probability distribution *P*(*x*) of *x* in each window of the umbrella sampling for $(GKDSKD)_7$.**

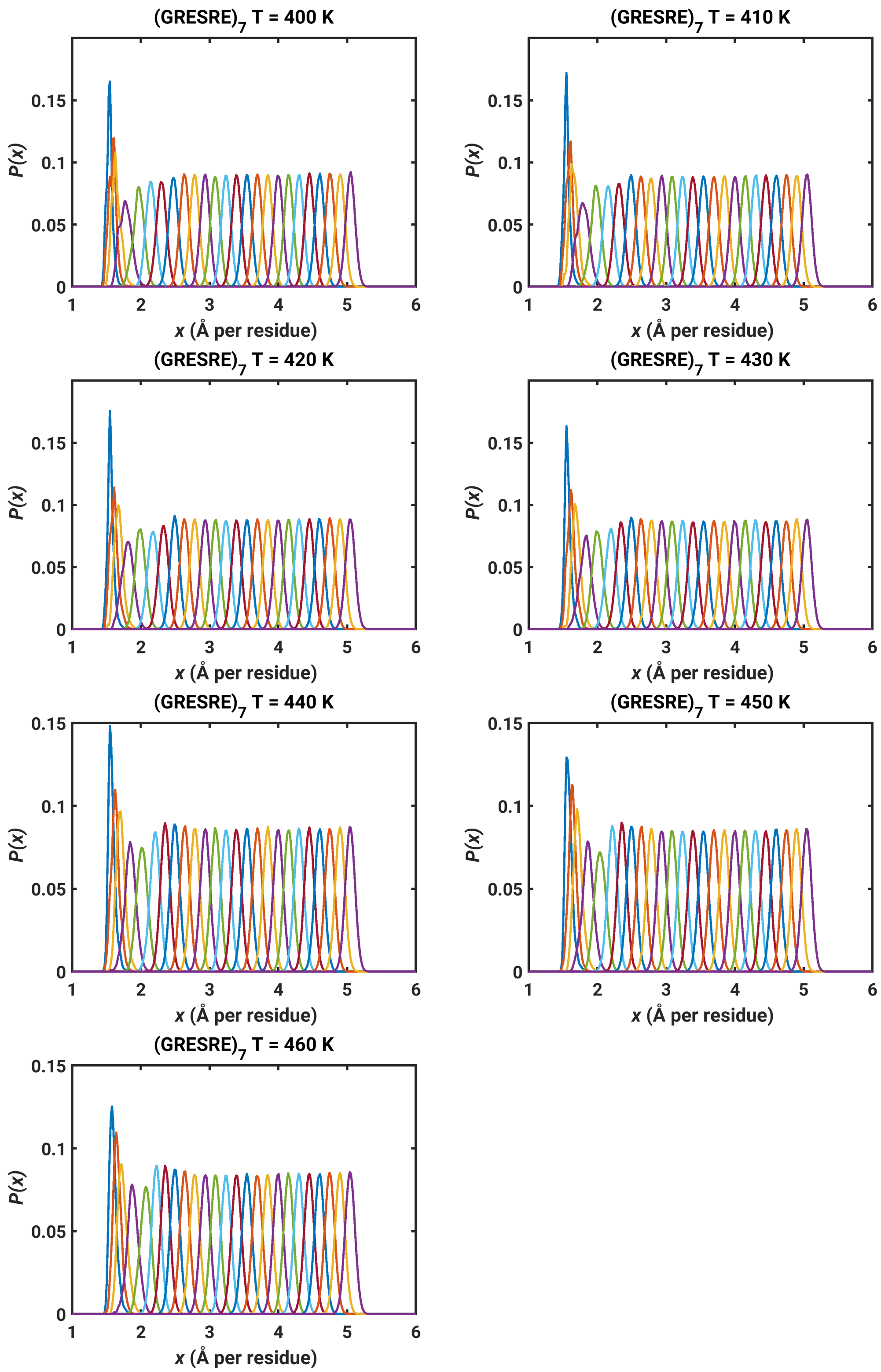


**Fig. S22. Probability distribution *P*(*x*) of *x* in each window of the umbrella sampling for (GRESRE)$_7$.**

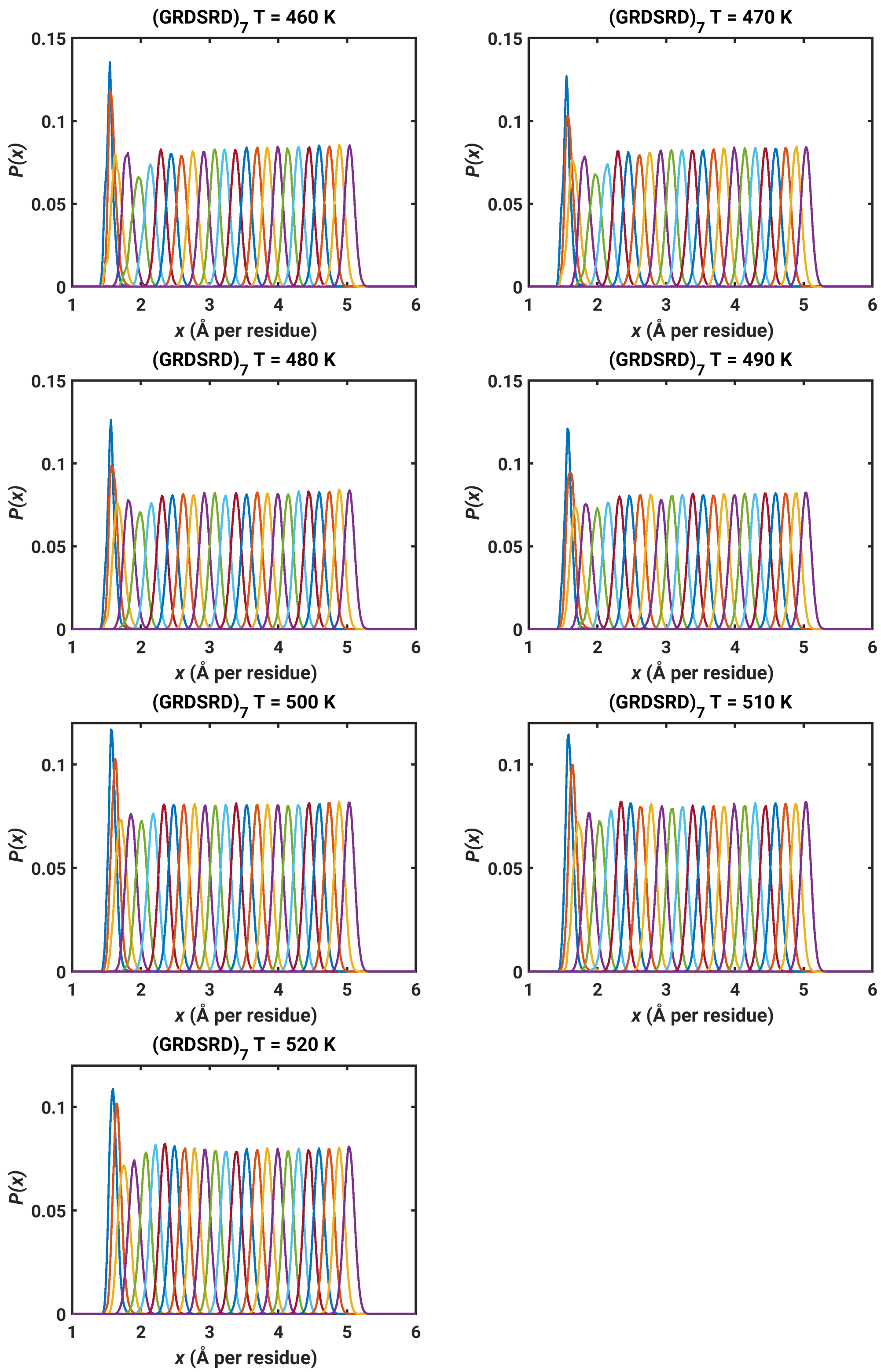


**Fig. S23. Probability distribution $P(x)$ of $x$ in each window of the umbrella sampling for $(GRDSRD)_7$.**

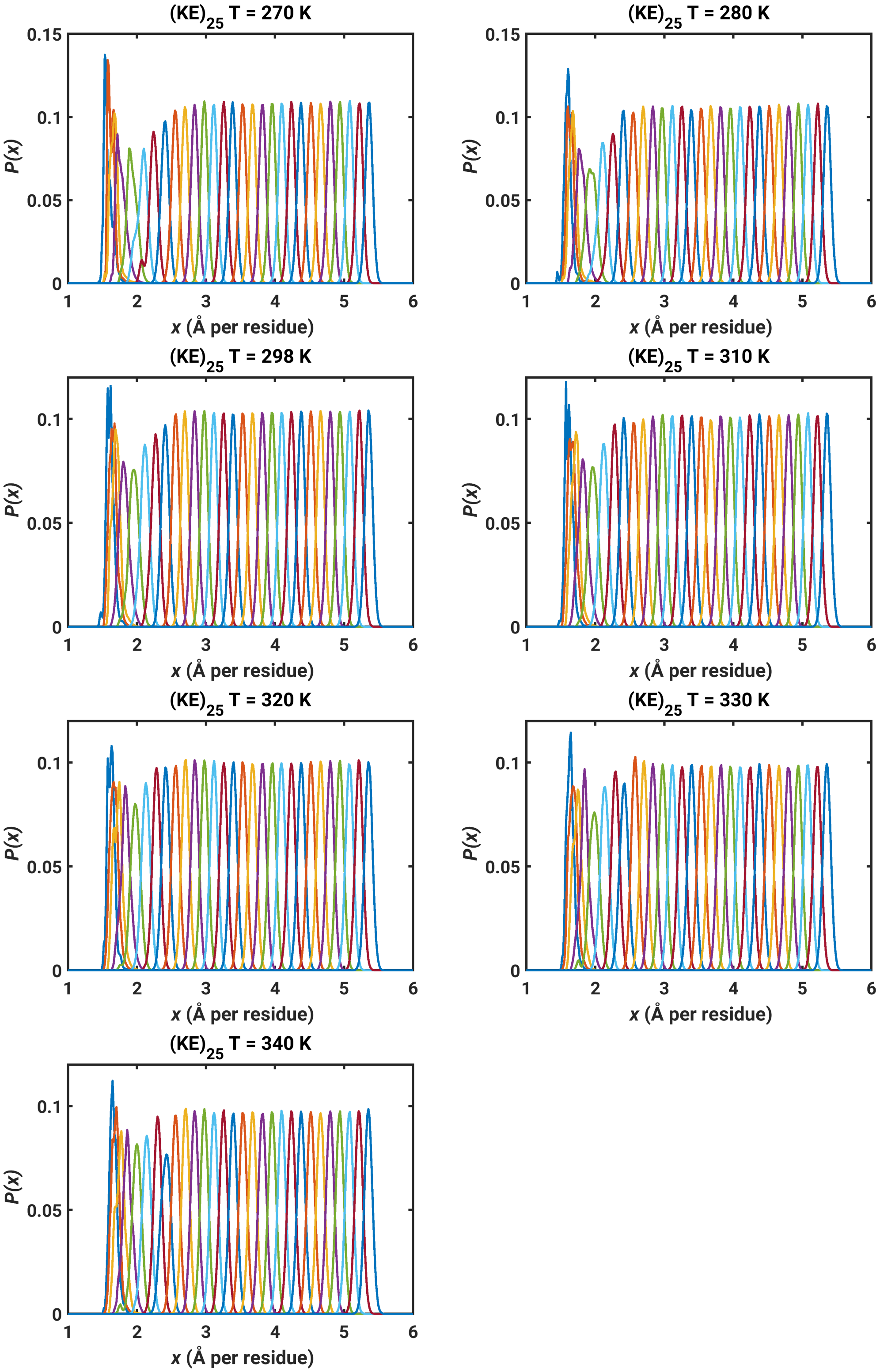


**Fig. S24. Probability distribution $P(x)$ of $x$ in each window of the umbrella sampling for $(KE)_{25}$.**

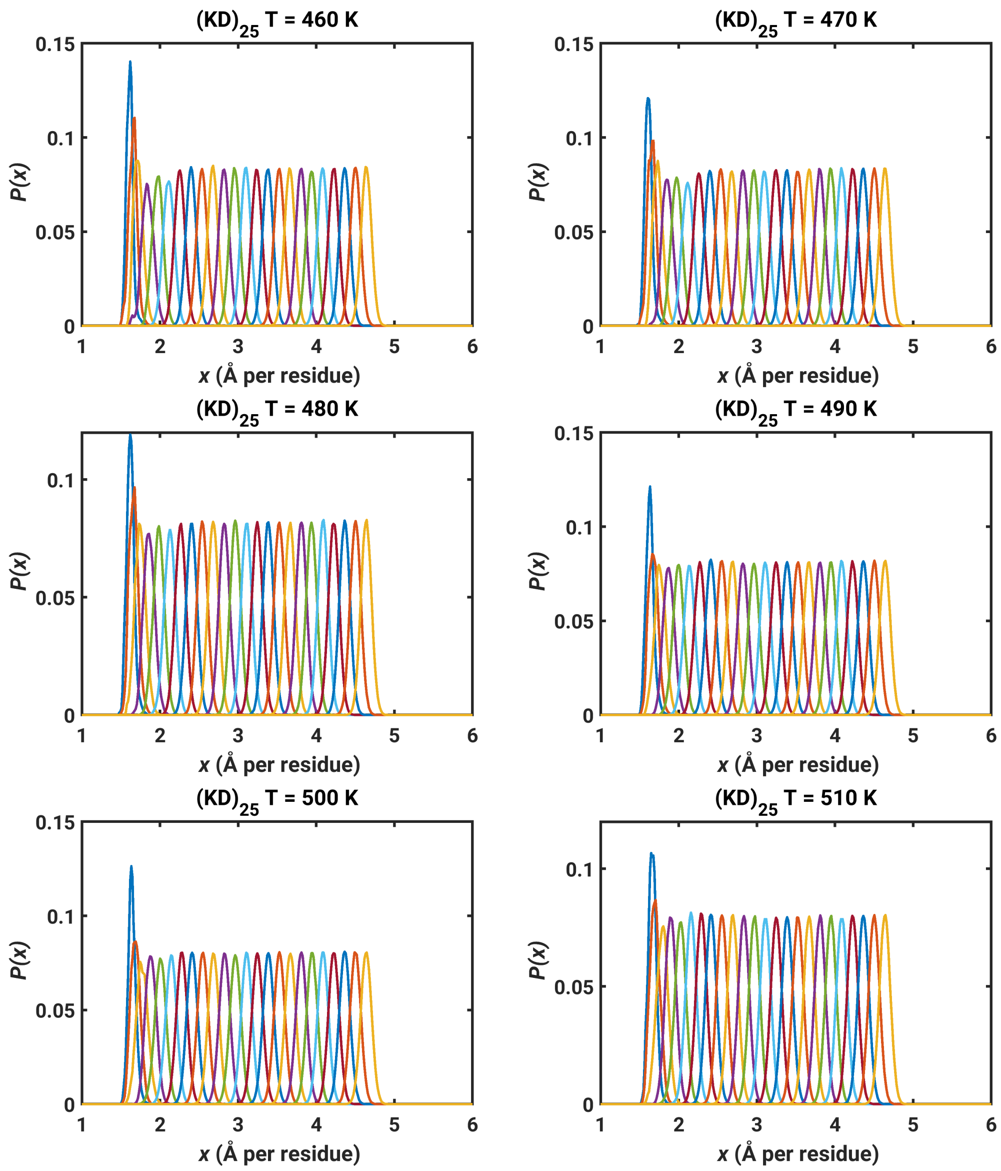


**Fig. S25. Probability distribution *P*(*x*) of *x* in each window of the umbrella sampling for (KD)$_{25}$.**

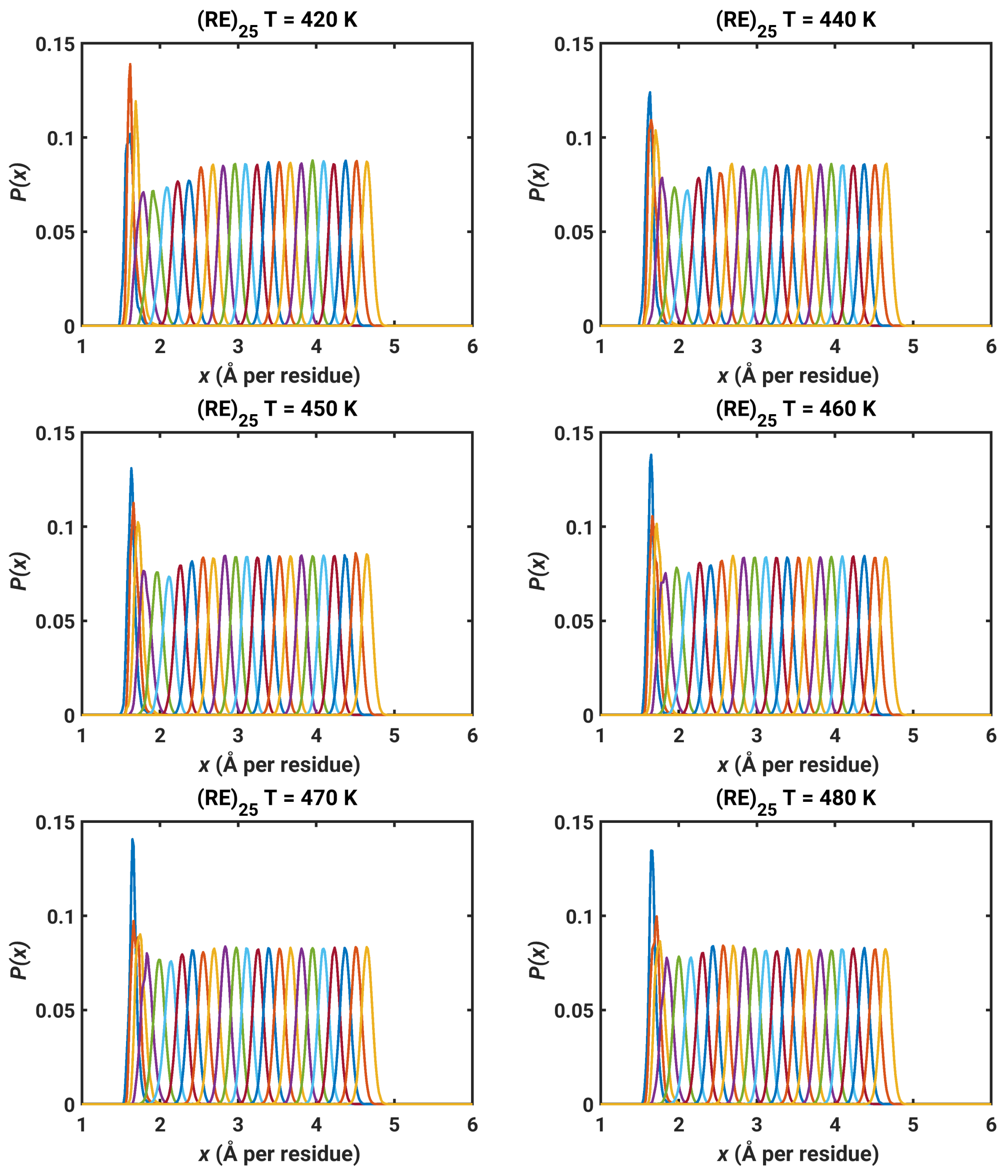


**Fig. S26. Probability distribution *P*(*x*) of *x* in each window of the umbrella sampling for $(RE)_{25}$.**

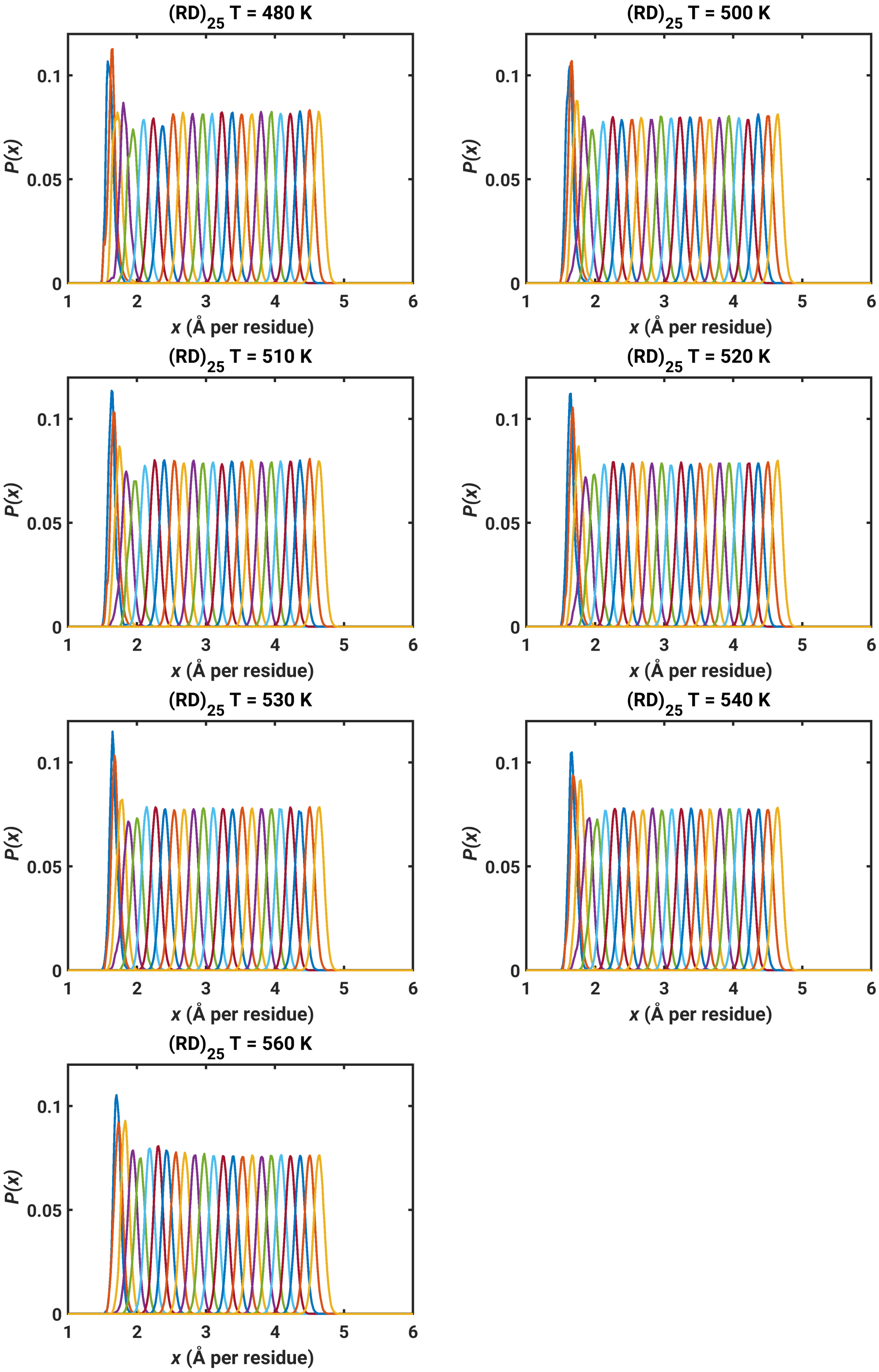


**Fig. S27. Probability distribution $P(x)$ of $x$ in each window of the umbrella sampling for $(RD)_{25}$.**

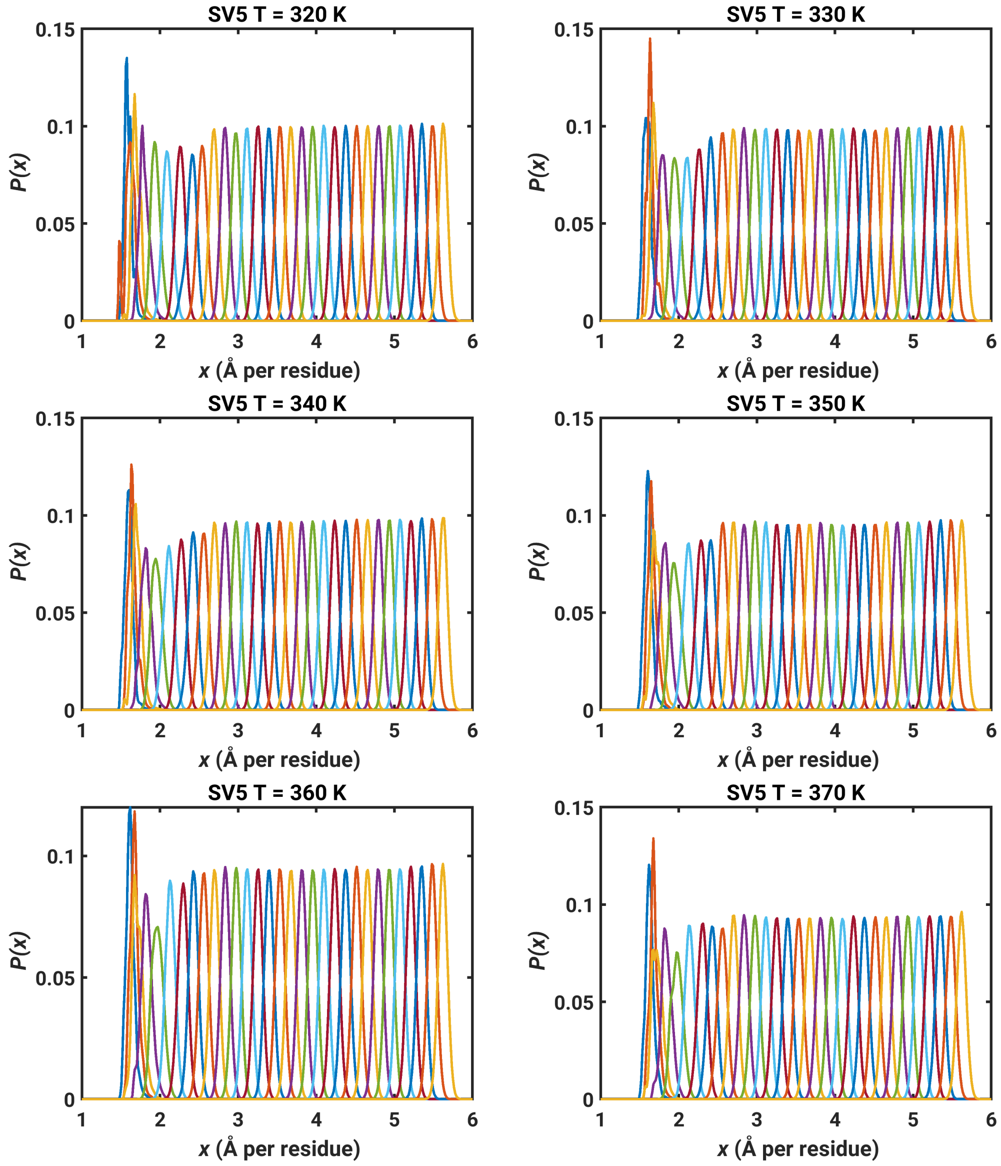


**Fig. S28. Probability distribution *P*(*x*) of *x* in each window of the umbrella sampling for sv5.**

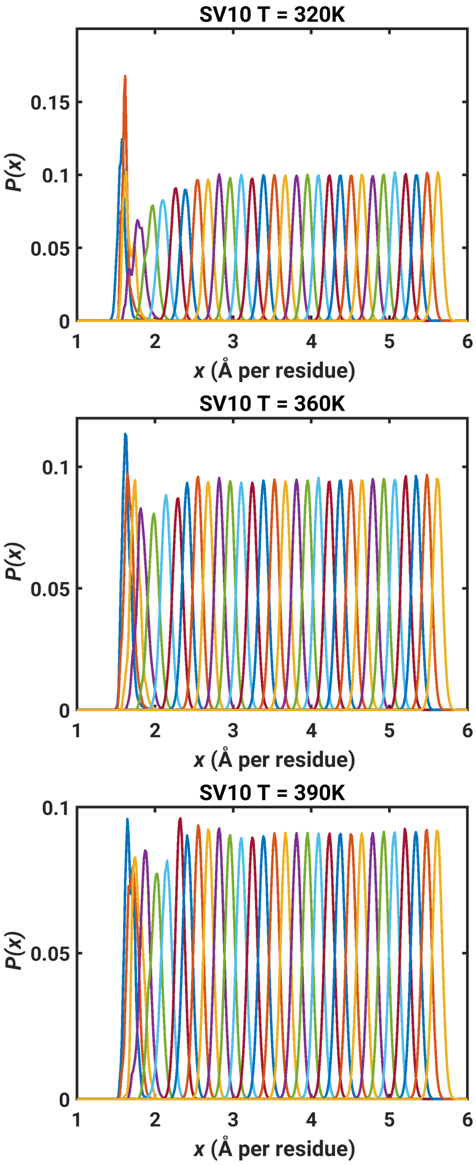


**Fig. S29. Probability distribution *P*(*x*) of *x* in each window of the umbrella sampling for sv10.**
